# Pneumococcus uses COMMD2 to alter host cellular immunity

**DOI:** 10.1101/2022.04.08.487599

**Authors:** Michael G. Connor, Lisa Sanchez, Christine Chevalier, Tiphaine M. N. Camarasa, Filipe Carvalho, Matthew J.G. Eldridge, Thibault Chaze, Mariette Matondo, Esma Karkeni, Sara Dufour, Francis Impens, Sebastian Baumgarten, Jost Enninga, Melanie A. Hamon

**Affiliations:** Chromatin and Infection, Institut Pasteur, Paris, France; Dynamics of Host–Pathogen Interactions Unit, Institut Pasteur, & UMR CNRS, Paris, France; Institut MICALIS (UMR 1319) INRAE, AgroParisTech, Université Paris-Saclay; Institut Pasteur, Université de Paris Cité, CNRS UAR2024, Proteomics Platform, Mass Spectrometry for Biology Unit, 75015 Paris, France; Single Cell Biomarkers UTechS, Institut Pasteur, Université Paris Cité, Paris, France; VIB-UGent Center for Medical Biotechnology, VIB Proteomics Core, VIB, Department for Biomolecular Medicine, Ghent University, Ghent, Belgium; Institut Pasteur, Université Paris Cité, Parasite RNA Biology, F-75015 Paris, France

## Abstract

NF-κB driven cellular immunity is essential for both pro- and anti-inflammatory responses to microbes, which makes it one of the most frequently targeted pathways by bacteria during pathogenesis. How NF-κB tunes the epithelial response to *Streptococcus pneumoniae* across the spectrum of commensal to pathogenic outcomes is not fully understood. In this study, we compare a commensal-like 6B ST90 strain to an invasive TIGR4 isolate and demonstrate, through comparative mass spectrometry of the p65 interactome, TIGR4 challenge triggers a novel interaction of COMMD2 with p65 and p62. Mechanistically, we show this complex mediates export of p65 for degradation and COMMD2 is necessary for altering host cellular immunity. With these results, we reveal for the first time a new bacterial pathogenesis mechanism to repress host inflammatory response though COMMD2 and p65 degradation while presenting a paradigm for diverging NF-κB responses to pneumococcus.

## Introduction

Airway epithelial cells, which line the entirety of the human respiratory tract, play a key role in both driving and shaping the host inflammatory and homeostatic response(s) to both pathogenic and commensal microbes ^1–3^. Indeed, defining molecular processes of bacterial-host crosstalk at this interface has emerged as a key determinant shaping how bacterial pathobionts escape from asymptomatic carriage within the human upper respiratory tract (URT) to become pathological agents causing symptomatic disease. In fact, for human restricted pathobionts, such as *Streptococcus pneumoniae* (the pneumococcus), *Haemophilus influenzae*, *Moraxella catarrhalis*, and *Staphylococcus aureus,* colonization of the human URT is critical for maintenance of their natural reservoir and a prerequisite for potentially lethal disease ^4,5^. Yet mechanistic understanding of the cellular processes involved in compartmentalizing commensal-like versus pathogenic host outcomes to these pathobionts remains unclear.

We previously showed that isolates of *Streptococcus pneumoniae* drive differential host responses by fine tuning airway epithelia cellular immunity ^6–12^. A commensal-like serotype 6B ST90 CC156 lineage F isolate, in contrast to an invasive favoring TIGR4 isolate, triggered a unique NF-κB signature through the chromatin remodeler KDM6B, that was required for host mediated confinement to the upper respiratory tract in a murine infection model. Surprisingly, these studies also revealed that the pathogenic *S. pneumoniae* strain TIGR4 largely suppressed NF-κB activation and downstream gene transcription ^6^. As the eukaryotic NF-κB family of transcriptional regulators are well documented for their potent ability to drive both pro- and anti-inflammatory cellular immune responses during microbe-host interaction, we sought to better define this dichotomy ^13–15^.

To date, a large majority of bacterial virulence phenotypes either co-opt NF-κB repressors or directly target NF-κB pathway components using virulence factors (general review ^15,16^). Generally, upon sensing of inflammatory molecules, such as cytokines (IL-1β or TNFα), pathogen-associated molecular patterns (PAMPs; i.e. lipopolysaccharide), or danger-associated molecular patterns (DAMPs; i.e. IL-1α or nuclear protein HMGB1), NF-κB subunits are rapidly activated by phosphorylation on serine residues (S536 and S276). Simultaneously, NF-κB dimers are released from their inhibitory IkB proteins and translocated to the nucleus for additional modification. Ultimately, this process leads to the binding of activated dimers to cognate NF-κB DNA motifs, thereby inducing NF-κB dependent gene transcription. NF-κB activation is tightly controlled for precise and rapid induction, but also for prompt repression. NF-κB dimers can be repressed through extraction, sequestration, and degradation from within the nucleus, while in parallel blocking cytoplasmic activation and promoting transcription of negative regulators ^17,18,20,22–26^. However, in contrast to the myriad of studies on activators only a few negative regulators of NF-κB and their pathways have been documented.

COMMD (<u>co</u>pper <u>m</u>etabolism gene <u>M</u>URR1 <u>d</u>omain)^27^ proteins are among the select few known negative regulators of NF-κB ^25,27–32^. There are ten members of the COMMD family, all of which, interact with NF-κB to regulate signaling. The best-studied architype member, COMMD1, upon stimulation by TNF will lead to extraction of p65 from chromatin, followed by ubiquitination and proteasomal degradation. This process functions independently of NF-κB nuclear translocation and IkBa, but through association with Cullin proteins, is able to terminate NF-κB signaling ^25,27,29–35^. For the other COMMD proteins, however, neither their functional activity, their mechanism of NF-κB repression, or their interacting partners, outside of cullins, are known.

Here, we demonstrate that a pathogenic TIGR4 pneumococcal strain ^6^, in contrast to a commensal-like 6B ST90 isolate ^6^, represses phosphorylation and activation of NF-κB p65. In fact, TIGR4 infection leads to specific degradation of p65 in airway epithelial cells, even upon stimulation with a strong inflammatory agonist, IL-1β. We performed an interactome of p65 and show that each pneumococcal strain interacts with diverging p65 interacting partners, revealing an original aggrephagy mechanism involving COMMD2 and p62. Furthermore, our studies on the understudied functionality of COMMD2 by RNAi demonstrated it is necessary in regulating host cellular immunity both basally and during pneumococcal challenge. Finally, deeper investigation of the COMMD2 interactome in comparison to the 6B ST90 isolate and under stimulation with the potent pro-inflammatory cytokine IL-1β revealed further specificity during TIGR4 challenge, with its COMMD2 interactome enriching for additional transcriptional and translational components. Altogether, we report a novel mechanism of pathogenesis to degrade p65 and repress the host response that is specifically induced by TIGR4.

## Results

### TIGR4 engages a divergent NF-κB p65 interactome in comparison to commensal-like 6B ST90

We previously showed that TIGR4, in comparison to 6B ST90 (hereafter 6B), induced a transcriptionally different NF-kB dependent response, which was largely associated with gene repression ^6^. To understand NF-κB p65 activation differences between the invasive TIGR4 and commensal-like 6B pneumococcal isolates we performed mass spectrometry of NF-κB p65 (Fig. 1A). An A549 GFP-p65 cell line was challenged with either TIGR4 or 6B and 2 hrs post-challenge GFP-p65 was immunoprecipitated with a matched A549 GFP alone control for mass spectrometry interactome analysis along with p65 posttranslational modifications (Fig. 1B; Sup. Table 1).

**Figure 1:**
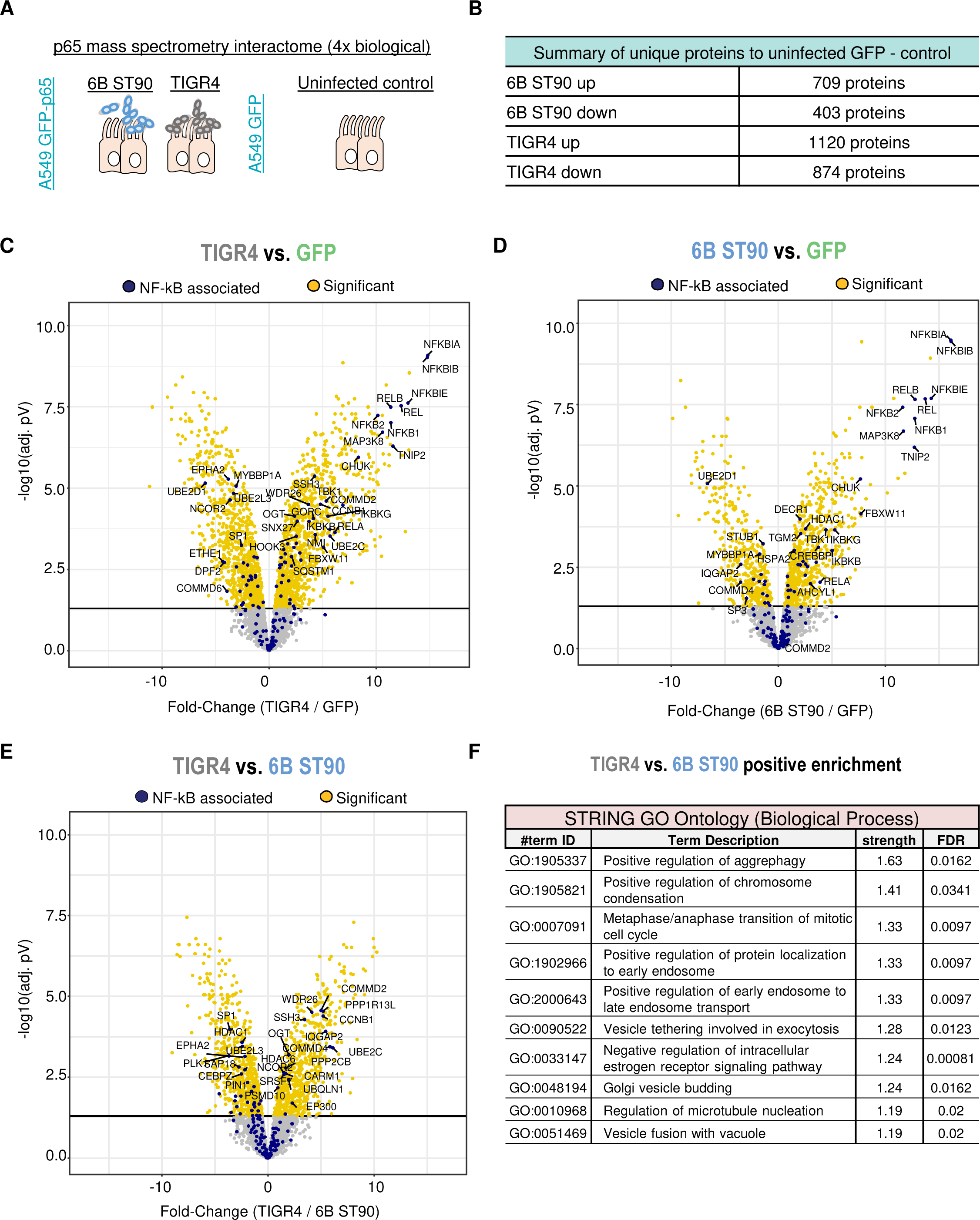
TIGR4 p65 interactome is divergent from 6B ST90 and enriches for aggrephagy. Mass-spectrometry interactome (n=4 biological replicates per condition) of immunoprecipitated GFP-p65 from a stable A549 GFP-p65 cell line (1×10^7^ cells total) 2 hrs post challenge with either 6B ST90 or TIGR4 (MOI 20). A) Experimental scheme of mass-spectrometry study. B) Tabular summary of NF-kB protein partners for TIGR4 and 6B in comparison to the GFP control. C-E) Volcano plots of TIGR4 vs. GFP (C), 6B ST90 vs. GFP (D), and TIGR4 vs. 6B ST90 (E). Known NF-kB p65 partners in blue and general significant targets in yellow. Lines represent −log10(pV) and fold-change (log2) cutoffs with targets of interested denoted. F) STRING GO Ontology analysis of significant TIGR4 targets with term strength and FDR. Full data in Sup. Table 1.

Analysis of the individual TIGR4 and 6B volcano plots in comparison to GFP alone showed specific enrichment for known NF-kB p65 associated partners (blue), reflecting the robustness of our A549 GFP-p65 cell line (Fig. 1C & D). GO ontology analysis of these individual p65 interactomes to GFP alone showed TIGR4 enriched for cellular trafficking components along with p62/SQSTM1 (Fig. 1C; Sup. Fig 1A), whereas the interactome of 6B reflected enrichment for classical NF-kB signaling components and signaling processes (Fig. 1D; Sup. Fig 1B). While the overall interactomes of TIGR4 and 6B appear similar, deeper investigation revealed key differences. This agrees with our previous findings demonstrating fine-tuning of NF-kB p65 signatures between TIGR4 and 6B was essential for commensal-like vs. invasive host responses ^6^.

However, to directly investigate any divergent p65 interactors between the TIGR4 and 6B datasets we compared the interactomes against each other (Fig. 1E). This comparison showed lower overall association of known NF-κB partners for TIGR4 alongside significant specific enrichment for COMMD proteins (2 & 4) and proteins associated with degradation/ubiquitin pathways (HDAC6, UBE2C…) by GO ontology (Fig. 1E & F; Sup. Table 1) ^36^.

Indeed, p62 and HDAC6 are classical mediators of aggrephagy, which is a form of selective autophagy to degrade protein aggregates ^36–38^. Interestingly, it has also been shown that abnormal NF-kB p65 activation may be a signal p62 mediated turnover ^39^. Moreover, COMMD proteins, such as COMMD2, are known to negatively regulate NF-kB family members through degradation ^27,29,30,34,35,40^. Of the two COMMD proteins found in our TIGR4 dataset, COMMD2 was previously shown, by ectopic expression, to reside in the cell cytosol and had a higher association with RelA and NF-κB1 ^27,35^. Therefore, our proteomic data suggested that p65 could be targeted for degradation through an aggrephagy pathway under TIGR4 infection conditions.

### TIGR4 differentially engages NF-κB p65 in comparison to commensal-like 6B ST90

Previously, we clearly showed commensal-like 6B pneumococcal strain activates p65 to drive a unique inflammatory signature in comparison to a disease causing TIGR4 strain ^6,41^. Interestingly, this was also seen in kinetic NF-kB dependent gene expression profiles when comparing TIGR4 to 6B and a positive control IL-1β ^6,41^. Given our previous findings and the mass spectrometry data showing enrichment for aggrephagy, we hypothesize that TIGR4 disrupted NF-κB activation. To directly test the NF-κB activation hypothesis, we challenged A549 cells with either TIGR4 or 6B ST90 alone, or in combination with IL-1β, a pro-inflammatory stimulus known to induce p65 phosphorylation of key serine residues 536 and 276 (review ^23^). Cells were collected 2hr post-challenge for immunoblotting to determine the total levels of p65 and phosphorylated p65. The ratio of phos-S536 to total p65 between TIGR4 and 6B (± IL-1β) against the IL-1β control showed decreased levels, whereas the ratio of phos-S276 to total p65 showed minimal differences (Fig. 2A – C). Curiously, for all TIGR4 challenge groups total level of p65 was significantly reduced (pV ≤ 0.01) compared to uninfected, IL-1β or 6B exposed cells (Fig. 2B). Thus, we compared phosphorylated p65 S536 and S276 to actin in order to determine whether TIGR4 challenge reduced their total cellular pool. Indeed, from this analysis it was clear that TIGR4 in comparison to 6B, reduced the total cellular amount of S536 p65 and not S276 (Sup. Fig. 2 A & B). Moreover, through cross-comparison between TIGR4 and 6B (± IL-1β) a trend emerged that either TIGR4 condition reduced total p65 S536 or S276 amounts, with the most substantial difference being on levels of p65 S536 (Sup. Fig. 2 A & B). Next, we tested whether TIGR4 challenge also lowered the level of p65 S536 phosphorylation in primary human nasal epithelial cells, since *S. pneumoniae* is an opportunistic respiratory pathogen that primarily colonizes first the upper airway epithelial cells. Our results showed a comparable trend for TIGR4 (± IL-1β) in comparison to uninfected, IL-1β or 6B. Importantly in TIGR4 lysates the total levels of p65, S536 and S276 were decreased in comparison to uninfected, IL-1β and 6B groups (Sup. Fig. 2C). Altogether, these data clearly indicated that TIGR4 challenge reduced total amounts of p65, phos-S536 and phos-S276, which primarily affected the ratio of phos-S536 to total p65, and aligned well with the interactome data.

**Figure 2:**
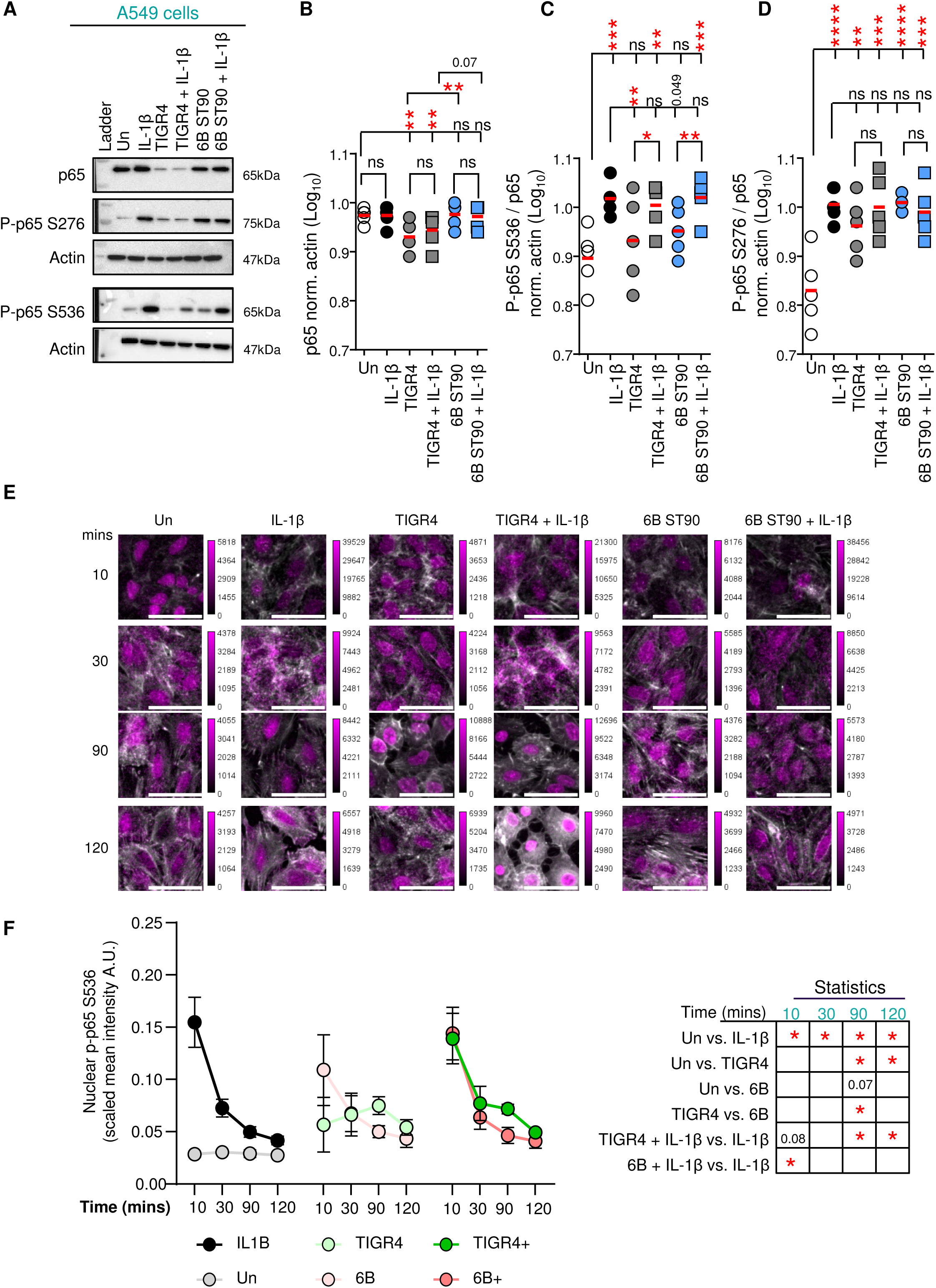
TIGR4 engages NF-kB differently than commensal-like 6B ST90. Immunoblot of A549 human airway epithelial cells 2 hrs post-challenge with either IL-1β (10 ng/ml), TIGR4 (MOI 20) or 6B ST90 (MOI 20) (± IL-1β; 10 ng/ml). Whole cell lysates probed for p65, phosphorylated p65 at Serine 276, phosphorylated p65 at Serine 536 or Actin. A) Representative image of immunoblot. Actin normalized ratiometric B) phosphorylated p65 at Serine 5366 to total p65, C) phosphorylated p65 at Serine 276 to total p65, and D) total p65. Dot blot with mean (red line). One-way ANOVA with Dunnett’s multiple comparison post-hoc test for B & C. One-way ANOVA with Tukey’s multiple comparison post-hoc test for D. Intergroup comparison paired T-Test for B-D. **P ≤ 0.01, ***P ≤ 0.001, ****P ≤ 0.0001. Full blots provided in Supplementary Information 1. E) Representative BioTek objective focused Zstack (10 µm) immunofluorescence microscopy of paraformaldehyde fixed A549 cells 2 h post-challenge with either IL-1β (10 ng/ml) or TIGR4 (± IL-1β 10 ng/ml; MOI 20) stained for phosphorylated p65 S536 (magenta) and actin (phalloidin; gray). Scale bar = 50µm. F) Quantification of scaled (x16bit) mean nuclear intensity of phosphorylated p65 S536. Dot blot of mean ± SEM with connected line over time (n = 3 biological replicates; counts in Sup. Table 4). Comparisons are Paired T-Test at each time indicated. * = significant; pV reported in Sup. Table 4.

As p65 phos-S536 is a marker reflecting p65 nuclear translocation (review ^23^), our observations that TIGR4 challenge resulted in significant difference in the S536 / p65 ratio raised the possibility that TIGR4 challenge induced differential nuclear localization kinetics in comparison to 6B. To assess this we completed time course microscopy of A549 cells to evaluate nuclear S536 levels by immunofluorescence, as this reflects p65 translocation post liberation from the cytoplasmic NEMO complex (review ^23^). For this, we segmented on cellular nuclei (Draq5) and quantified nuclear p65 S536 intensity (Fig. 2E & F). Our kinetics show nuclear S536 profile between TIGR4 and 6B differs. TIGR4 leads to a slow and modest increase starting at 10 minutes and peaks at 90 mins before coming back down. In contrast 6B kinetics mirror the positive control, IL-1β, with a rapid increase by 10 minutes followed by a gradual decline (Fig. 2E & F). When A549 are challenged with either TIGR4 or 6B in combination with IL-1β, the levels of nuclear S536 increase rapidly for both, similarly to IL-1β alone. These data supported differences observed by immunoblot, whilst suggesting that TIGR4 (± IL-1β) was triggering aberrant and low level p65 activation in comparison to IL-1β and 6B.

Since we previously showed that TIGR4 alone, in comparison to 6B and IL-1β, drove minimal NF-kB p65 dependent gene expression kinetically ^6,41^, we tested if IL-1β at the time of challenge could restore expression profiles to TIGR4. Total RNA was periodically collected from A549 cells out to 120 minutes post challenge with TIGR4 (± IL-1β) and IL-1β alone. Relative expression was determined for CFS2 and PTGS2 (COX-2) against uninfected/untreated controls at each time point. TIGR4 in comparison with IL-1β alone did not lead to activation of any of the genes out to 2h post challenge, which aligns with our previous findings ^6^ (Sup. Fig. 2D & E). Furthermore, under conditions where IL-1β was added during TIGR4 challenge, there was both a delay and a repression of these transcripts in comparison to IL-1β alone.

Transcriptional activation by p65 requires its binding to cognate kappa-binding sites at the chromatin level. Therefore, we evaluated levels of chromatin bound p65 at the locus of the NF-κB dependent gene *PTGS2* and *CSF2* as these two genes had striking transcriptional profiles. Chromatin was collected from A549 cells 2 hrs post-challenge with TIGR4 (± IL-1β) and the recovery of p65 quantified against uninfected and IL-1β controls by ChIP-qPCR targeting kappa-binding sites upstream of the *PTGS2* and *CSF2* transcriptional start site (Sup. Fig. 2F - H). For both genes TIGR4 challenged cells resulted in less than 5% recovery of p65. This stands in contrast to the three-fold higher p65 recovery in IL-1β alone (Sup. Fig. 2G & H). Interestingly, in the TIGR4 + IL-1β condition p65 recovery was comparable to IL-1β alone (Sup. Fig. 2G & H). Therefore, the lack of p65 driven transcription under TIGR4 challenge is intrinsically due to the absence of p65 at the chromatin. Altogether, these results, in combination with our previous findings, show that TIGR4, in contrast to the IL-1β, is driving dysfunctional p65 signaling ^6,41^.

### Blocking autophagy increases cytoplasmic COMMD2, p62 and p65 levels

To study the localization of COMMD2, we generated a COMMD2 cell line. Strikingly, upon TIGR4 challenge of the GFP-COMMD2, we observed by microscopy that TIGR4 drove COMMD2 localization to the nucleus. COMMD2 is an otherwise cytoplasmic protein containing two nuclear export signals ^34,35,43^. We confirmed this translocation by immunoblotting GFP-COMMD2 cell fractionations obtained from cells challenged with TIGR4 (± IL-1β) alongside from uninfected and IL-1β controls (Sup. Fig. 3A – C). Cells treated with IL-1β or uninfected had 20% COMMD2 in the nucleus, in contrast to that of TIGR4 (± IL-1β) which showed 80% of COMMD2 was nuclear (Sup. Fig. 3B). Finally, we tested if the commensal-like strain 6B ST90 could also induce COMMD2 nuclear localization. Immunoblotting of cell fractions of the GFP-COMMD2 we show 6B ST90 was incapable of triggering nuclear localization of COMMD2 (Sup. Fig. 3C). Thus, these data demonstrate TIGR4 specifically triggers COMMD2 nuclear localization.

**Figure 3:**
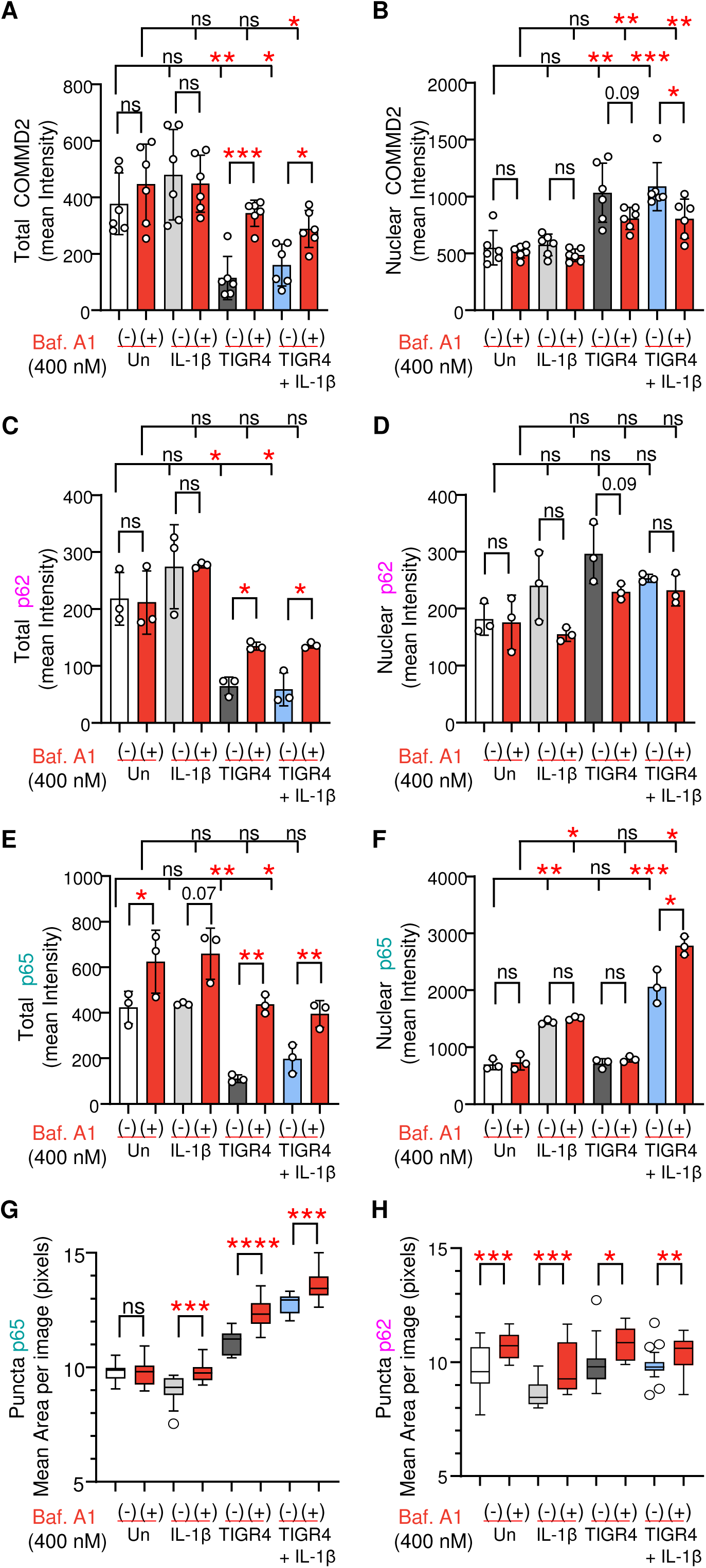
Blocking aggrephagy increases total COMMD2, p62 and p65. Immunofluorescence confocal microscopy of stable A549 GFP-COMMD2 cells pre-treated with Bafilomycin A1 (400nM) or DMSO for 3 h prior to 2 h post-challenge with either IL-1β (10 ng/ml) or TIGR4 (± IL-1β 10 ng/ml; MOI 20). Representative images Sup. Fig. 4 & 5. Bar graph of mean ± STD (n = 3 biological replicates for p65 and p62; n = 6 biological replicates for COMMD2; counts in Sup. Table 4). A & B) Quantified levels of total or nuclear COMMD2. C & D) Quantified levels of total or nuclear p62. E & F) Quantified levels of total or nuclear p65. G &) Quantified area (pixels) of total p65 puncta or p62 puncta (n = 3 biological replicates for p65 and p62). For G & H Tukey box and whisker plot with defined box boundaries being the upper and lower interquartile range (IQR), whiskers’ (fences) being ± 1.5 times IQR and the median depicted by the middle solid line. Dots represent outliers. One-way ANOVA with Tukey’s multiple comparison post-hoc test for A - H. **P ≤ 0.01, ***P ≤ 0.001, ****P ≤ 0.0001.

Using the COMMD2 cell line, we tested the effect of blocking aggrephagy with Bafilomycin A1 on the levels of p65, p62 and COMMD2, 2 hrs post-challenge with either IL-1β or TIGR4 (± IL-1β; MOI 20) using confocal microscopy (Fig. 3A – F; rep. images Sup. Fig. 4 p65 & Sup. Fig. 5 p62). Baf. A1 treatment had minimal effect on total or nuclear COMMD2 for uninfected and IL-1β, whereas TIGR4 (± IL-1β) stimulation significantly (pV ≤ 0.001) increased total COMMD2 (Fig. 3 A &B). Interestingly, there was a slight inverse decrease in TIGR4 (± IL-1β) nuclear COMMD2 – strongly supporting a role for aggrephagy in cytoplasmic turnover. Similarly to COMMD2, TIGR4 (± IL-1β) challenged cells treated with Baf. A1 displayed an increase in p62, and restored p65 levels comparable to uninfected controls (Fig. 3C - F). Moreover, in Baf. A1 treated cells challenged with TIGR4 the area of p65 and p62 puncta increased, which is a conventional phenotype associated with blocked autophagosomes (Fig. 3G & H; rep. images Sup. Fig. 4 p65 & Sup. Fig. 5 p62).

**Figure 4:**
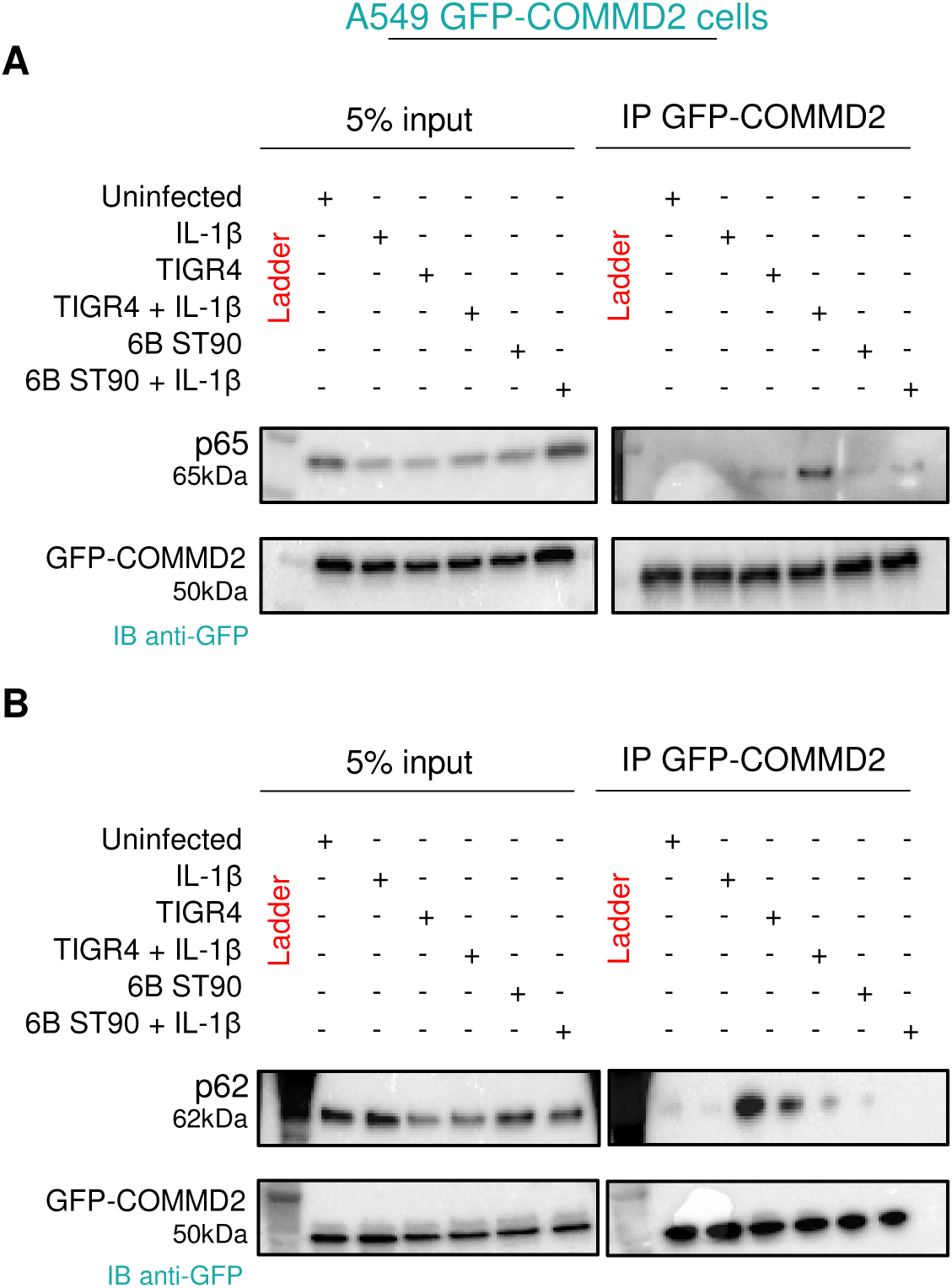
TIGR4 challenge induces formation of a COMMD2 – p65 – p62 complex. GFP-Trap agarose immunoprecipitants were collected from a stable A549 GFP-COMMD2 cell line 2 hrs post-challenge with either IL-1β (10 ng/ml) or TIGR4 (MOI 20) or 6B ST90 (MOI 20) (± IL-1β; 10 ng/ml). A & B) Representative immunoblot from 3 biological replicates of GFP-COMMD2 immunoprecipitation lysates (input & IP) probed for p65, p62 or GFP for COMMD2. Full blots provided in Supplementary Information 1.

**Figure 5:**
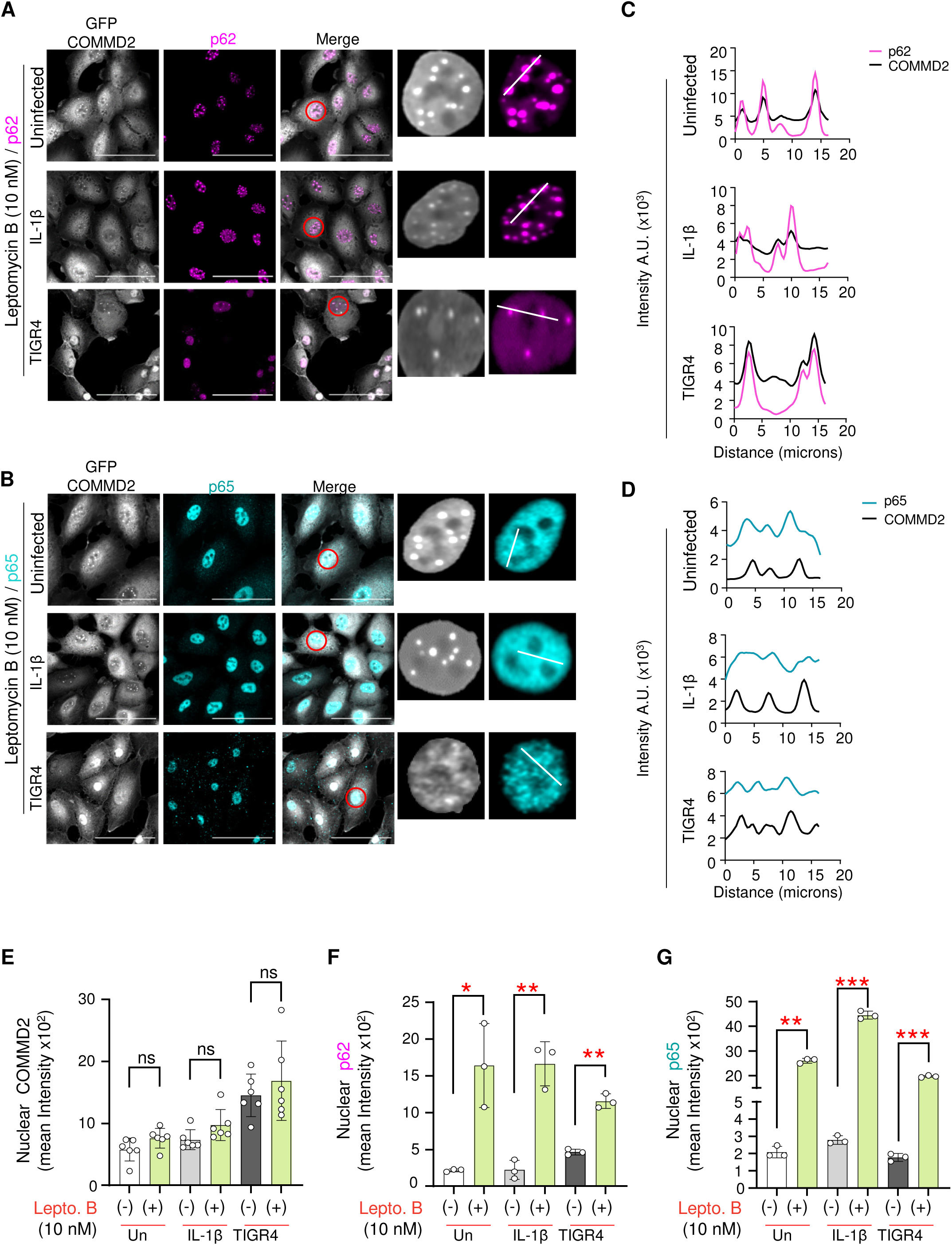
COMMD2-p65-p62 is exported from nucleus through CRM1. Representative immunofluorescence confocal microscopy of stable A549 GFP-COMMD2 pretreated for 3 hrs with Leptomycin B (10 nM) prior to 2 hr challenge with either IL-1β (10 ng/ml) or TIGR4 (MOI 20). Paraformaldehyde fixed cells stained for A) p62 (magenta), or B) p65 (cyan) against GFP-COMMD2 (gray). Scale bar = 100 µm or 10 µm for single nuclei inserts. C & D) Co-association of intensity peaks across a 20 µm segment (white line) for p62 (magenta) or p65 (cyan) to COMMD2 (black). E-G) Quantified nuclear intensity of COMMD2 (E), p62 (F), p65 (G) from untreated uninfected, IL-1β or TIGR4 conditions to Leptomycin B treatment. Graphed as mean ± STD with dots representing biological replicates. Intergroup comparisons paired T.Test. *P ≤ 0.05, **P ≤ 0.01, ****P ≤ 0.0001.

Overall, the combination of these data clearly shows TIGR4 challenge of airway epithelial cells specifically led to COMMD2 nuclear localization and aggrephagy, a largely cytoplasmic process, contributing to reduce the levels of p65 and COMMD2 within the cell.

### COMMD2 is associated with both p65 and p62

To further test the aggrephagy hypothesis, we wanted to determine whether COMMD2, p65 and p62 formed a complex; thus, a plausible explanation why blocking aggrephagy with Baf. A1 would increase all three of these components. Currently, the only commercial COMMD2 antibody has limited suitability for immunoprecipitation or immunofluorescence, thus, we used the GFP-COMMD2 cell line. For this we challenged cells with either 6B ST90 (± IL-1β), TIGR4 (± IL-1β) or IL-1β controls for 2h and performed immunoprecipitation of GFP-COMMD2 from cell lysates. Samples were probed for p65 or p62 to detect interaction with COMMD2. Our results show that only under TIGR4 challenge conditions do p65 and p62 interact with COMMD2 (Fig. 4A & B). Furthermore, upon addition of IL-1β, p65 interacts with COMMD2 to even higher levels.

To evaluate complex formation in primary cells, we collected lysates from primary nasal cells challenged with either TIGR4 or 6B, as well as from uninfected controls. We performed p65 immunoprecipitation and tested whether TIGR4 could drive an endogenous native interaction of p65 with COMMD2 (Sup. Fig. 6A). Samples were probed for p65 and COMMD2 prior to being revealed with the VeriBlot secondary HRP antibody. This secondary HRP antibody does not recognize denatured heavy and light chain IgG, which would otherwise occur closely to our targets potentially masking positive bands (65kda & 25kda). Our results showed a 4-fold increase in COMMD2 interaction with p65 under TIGR4 challenge (Sup. Fig. 6B). However, there was a significant reduction of the detected levels of COMMD2 and p65 under TIGR4 conditions in comparison to uninfected or 6B (Sup. Fig. 6C &D). This is consistent with results shown above and degradation of COMMD2 and p65 by aggrephagy. Interestingly, there was a weight shift of COMMD2 in the p65 IP, suggestive of posttranslational modification – known to occur on COMMD proteins. Therefore, COMMD2 is a new TIGR4 induced interacting partner of p65 and is a component of a p65-COMMD2-p62 complex.

**Figure 6:**
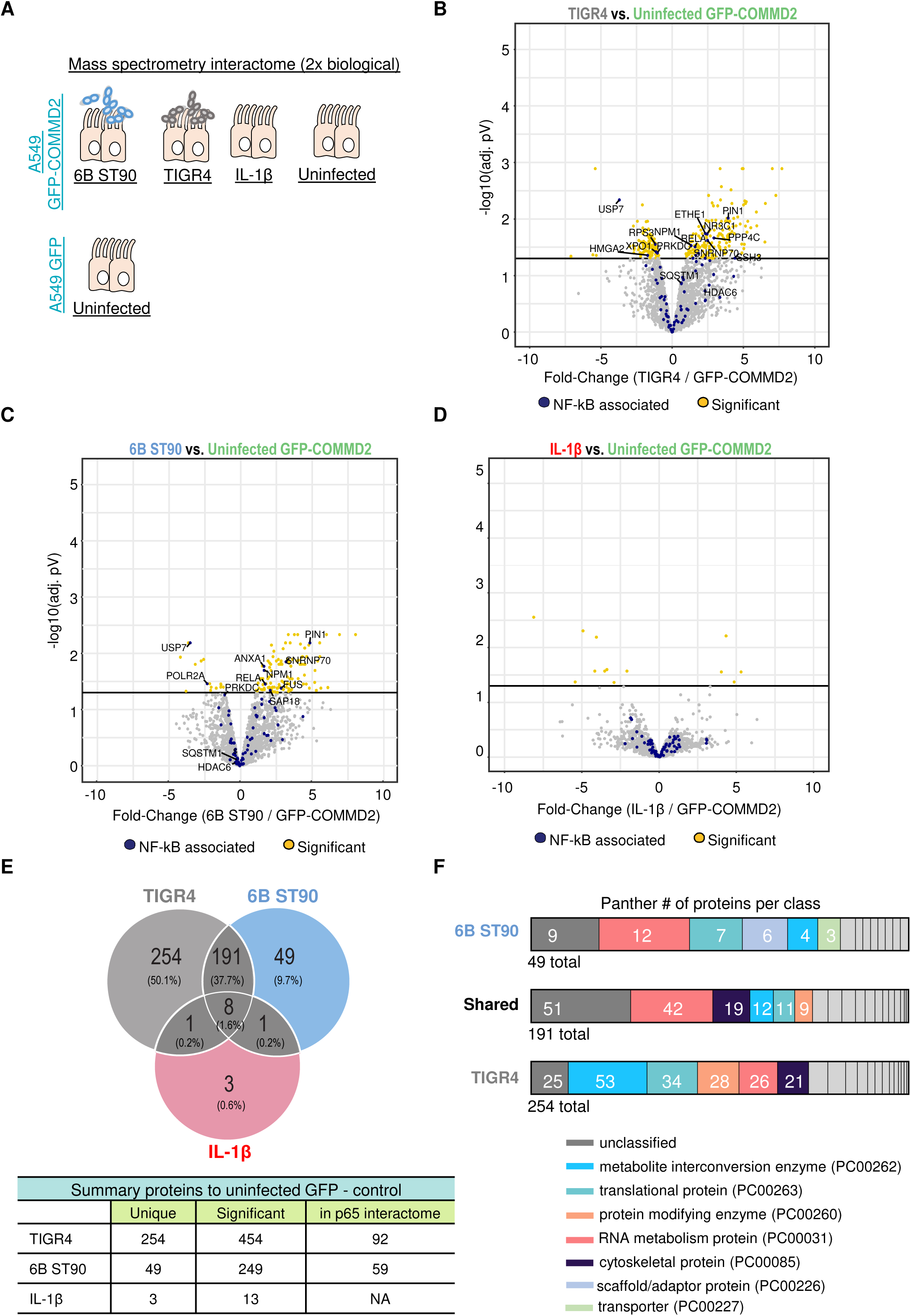
TIGR4 driven COMMD2 interactome is unique in comparison to 6B ST90 or IL-1β. Mass-spectrometry interactome (n=2 biological replicates per condition) of immunoprecipitated GFP-COMMD2 from a stable A549 GFP-COMMD2 cell line (1×10^7^ cells total) 2 hrs post challenge with either TIGR4 (MOI 20), 6B ST90 (MOI 20) or IL-1β (10 ng/ml). A) Experimental scheme of mass-spectrometry study. B – D) Volcano plots of TIGR4 vs. GFP-COMMD2 (B), 6B ST90 vs. GFP-COMMD2 (C), and IL-1β vs. GFP-COMMD2 (D). Identified interacting partners with known NF-kB p65 interaction in blue and significant targets in yellow. Lines represent −log10(pV) and fold-change cutoffs with targets of interested denoted. E) Venn diagram describing the numbers of unique and shared significant proteins and their total % from the mass-spectrometry dataset. Table breakdown of identified partners that were significant, of the significant targets that are unique, and of the significant targets shared with the p65 interactome. F) Panther classification analysis of protein class of identified COMMD2 interactors unique to TIGR4, shared between TIGR4 and 6B ST90, or unique to 6B ST90.

### COMMD2 exports p65 for lysosomal degradation

It is known that the architype COMMD family member, COMMD1, undergoes nucleocytoplasmic trafficking ^25,35,42^. Given that aggrephagy primarily occurs in the cellular cytosol, we reasoned that aberrantly phosphorylated p65 during TIGR4 challenge led to nuclear translocation of COMMD2, which in turn was involved with facilitating nuclear export of these p65 complexes.

Therefore, we tested whether COMMD2 was exported through CRM1 by using the Leptomycin B inhibitor ^43^. GFP-COMMD2 cells were treated with Leptomycin B (10 nM) prior to immunofluorescence imagining of p62 and p65. Strikingly, blocking nuclear export resulted in COMMD2, p62 and p65 now becoming localized to the nucleus even in uninfected and IL-1β treated cells. In fact, COMMD2 and p62 were detected within nuclear puncta (Fig. 5A) where p65 was located (Fig. 5B). Notably, our IL-1β positive pro-inflammatory stimulus control, known to drive nuclear translocation of p65, had a significant (pV ≤ 0.0001) increase in the nuclear level of COMMD2 and p62 in comparison to treated cells (Fig. 5C & D). These surprising data suggest that under nuclear export blockage, which occurs with Leptomycin B treatment ^43^, COMMD2 and p62 are naturally recruited to the nucleus at specific puncta through a defined process.

Interestingly, upon challenge with TIGR4, Leptomycin B treated cells displayed higher levels of COMMD2 and p62 accumulation in the nucleus than without treatment (Fig. 5A, C & D). Thus, upon inhibiting nuclear export of COMMD2 and p62, these proteins are no longer degraded upon infection and accumulate in the nucleus. Although COMMD2 and p62 puncta are observed upon addition of Leptomycin B, the substantial amount of p65 trapped within the nucleus of these cells rendered definitive scoring of puncta and colocalization for both difficult. Due to this, we measured a tangent line (20 µm) across observed local maxima intensity “spots” to overlay the traces of p62 and p65 with COMMD2 in uninfected, IL-1β and TIGR4 conditions demonstrating that indeed p62 and p65 associated with COMMD2 (Fig. 5C & D). Quantification of the total nuclear COMMD2 intensity however showed inhibition of CRM1 did not increase total levels in comparison to untreated across all groups (Fig. 5E). Thus, Leptomycin B inhibition led to concentrated accumulation of COMMD2 into puncta. In contrast, Leptomycin B inhibition increased both p62 and p65 nuclear levels in all conditions compared to untreated cells (Fig. 5F & G). Furthermore, under conditions of TIGR4 + IL-1β and Leptomycin B inhibition, we observed a significant (pV ≤ 0.0001) increase in p62 puncta compared to TIGR4 alone (Sup. Fig. 7A & B). These data show that TIGR4 challenge leads to an active CRM1 dependent export of p65 and the COMMD2 complex.

**Figure 7:**
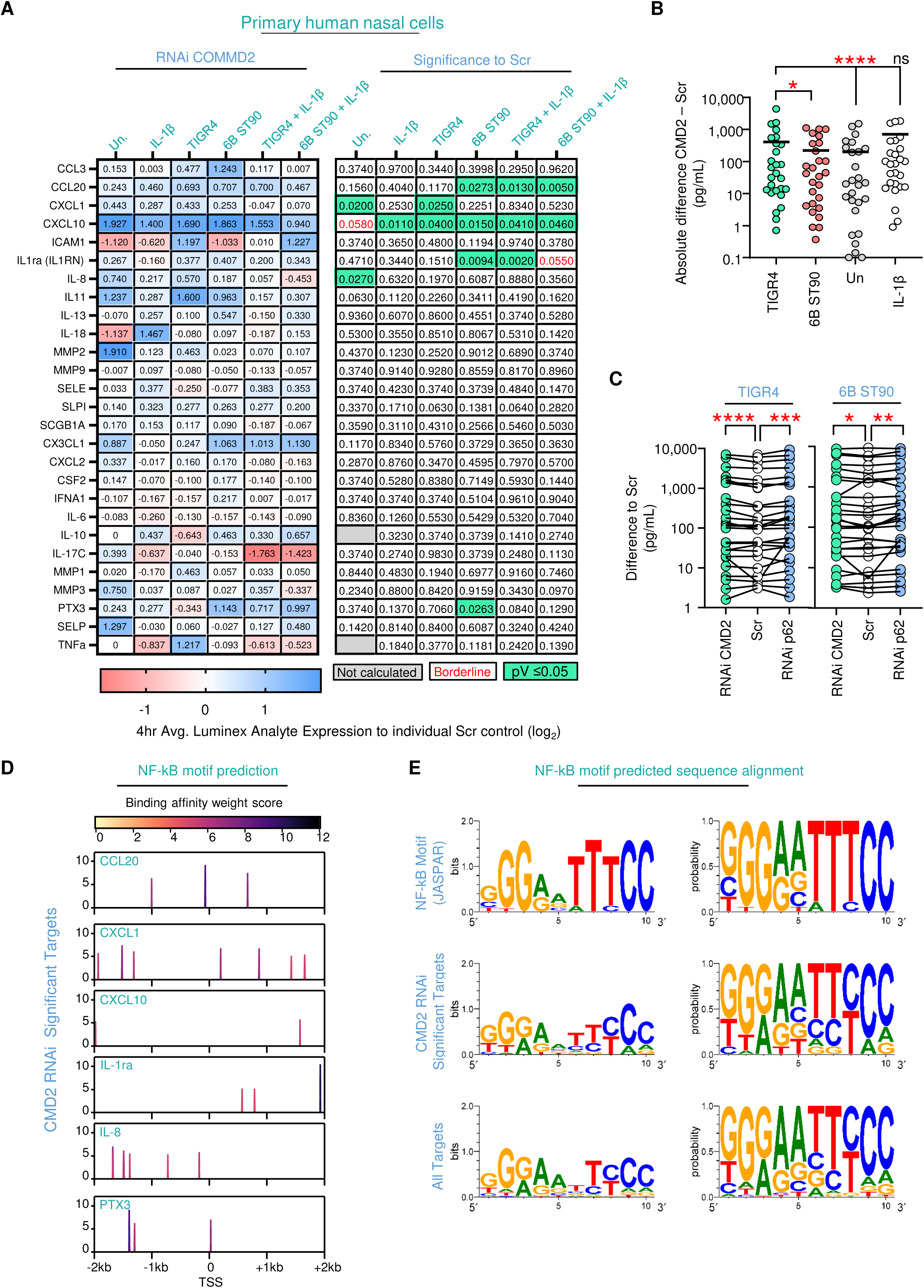
COMMD2 RNAi selectively elevates host analytes associated with cellular immunity. A) Secreted analytes (Luminex) at 4 hrs from RNAi COMMD2 primary nasal cells normalized to scrambled controls (n=3 biological replicates). Cells were challenged with either IL-1β (10 ng/ml) or TIGR4 (MOI 20) ± IL-1β (10 ng/ml). Individual p values shown in comparison to respected Scr control. Calculated by unpaired T. Test. B) Absolute difference of COMMD2 RNAi to scramble control for all analytes. Dot blot with mean (bars). Wilcoxon matched pairs signed rank test to TIGR4. ns = not significant, *P ≤ 0.05, ****P ≤ 0.0001. C) Difference of either RNAi COMMD2 or p62 against scramble control for TIGR4 and 6B ST90. Before-after line graph. Wilcoxon matched-pairs signed rank test to TIGR4. *P ≤ 0.05, **P ≤ 0.01,***P ≤ 0.001, ****P ≤ 0.0001. D) Transcription factor Affinity Prediction (TRAP) analysis ^47^ of predicted NF-kB binding motifs ± 2kb of indicated gene TSS. Scale bar represents predicted affinity score color coded. E) Weblogo ^48^ visual of nucleotide probability of CMD2 target genes in comparison to canonical JASPAR NF-kB motif and all targets. Full details of analysis in Sup. Table 4.

### The TIGR4 COMMD2 interactome is unique and enriched for NF-kB associated partners

Our results to this point clearly show the role of COMMD2–p65 during TIGR4 bacterial challenge is unique even with respect to IL-1β. However, to date knowledge of COMMD protein interacting partners and biological roles are largely centered around their interactions with NF-κB subunits and cellular processes of trafficking, proteasome degradation and transcription ^25,31,44,45^, with limited understanding of their potential mechanisms or impact during bacterial infection.

To deepen our investigation of TIGR4 induced COMMD2-p65 interaction and better define a biological role for it during bacterial challenge, we assessed the COMMD2 interactome by mass spectrometry (Fig. 6A-D). For this we immunoprecipitated GFP-COMMD2 from A549 cells 2 hrs post-challenge with either TIGR4, 6B ST90 or our positive inflammatory control IL-1β for comparison against uninfected. Pointedly in this mass spectrometry study we used the cross-comparison of GFP-COMMD2 against GFP alone to control for both non-specific COMMD2 complexes and artifacts which may be intrinsic to ectopic expression (Sup. Fig. 8A; Sup. Table 2). The results of this comparison identified known COMMD2 associations with other COMMD proteins, Commander and Retriever complexes (CCDC22, CDC93, VPS35L, COMMD6, COMMD2, COMMD1, COMMD4, COMMD8, COMMD9, COMMD3 & VPS29) ^46^, which further strengthens the robustness of our approach.

**Figure 8:**
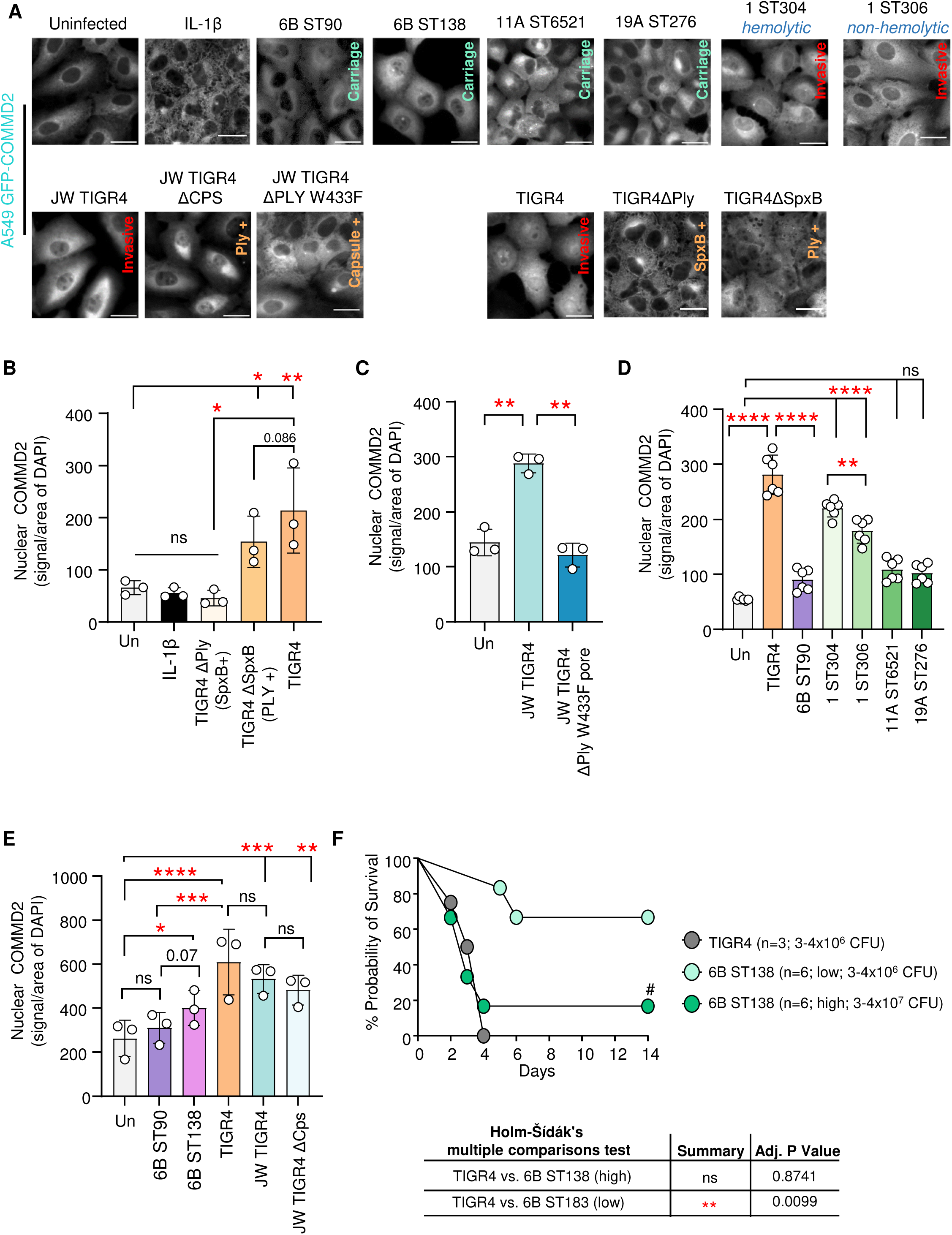
Invasive favoring pneumococcal isolates drive COMMD2 nuclear localization. A) Immunofluorescence deconvolution epifluorescence microscopy of A549 GFP-COMMD2 (gray) cells 2 hr post-challenge with either IL-1β (10 ng/ml), or MOI 20 for each bacterial strain 6B ST90, 6B ST138, 11A ST6521, 19A ST276, 1 ST304, 1 ST306, JW TIGR4 (JW parental wildtype), JW TIGR4 Δ*cps*, JW TIGR4 Δ*ply* W433F, TIGR4 (parental wildtype), TIGR4 *ΔspxB* or TIGR4 *Δply*. Relevant bacterial strain information for favoring carriage or invasive host states along with mutant genotypes. Scale bar = 50µm. B-E) Bar graph of mean ± STD quantification of nuclear GFP-COMMD2 normalized to the segmented nuclei using DAPI signal (omitted in representative images for clarity; B,D & E n = 3 biological replicates; C n = 6 biological replicates; counts in Sup. Table 4). One-way ANOVA with Tukey’s multiple comparison post-hoc test for A - H. **P ≤ 0.01, ***P ≤ 0.001, ****P ≤ 0.0001. F) C57B6 mice (8-9weeks) were intranasally challenged with TIGR4 (3 - 4×10^6^ CFU; n=3), 6B ST138 (3 - 4×10^6^ CFU; n=6), or 6B ST138 (3 - 4×10^7^ CFU; n=6). Survival monitored for 14 days post-infection. Kaplan-Meier percent survival curve with Holm-Šídák’s multiple comparisons test. **P ≤ 0.01. # = single animal survived.

In contrast to 6B and IL-1β volcano plots, TIGR4 prompted positive enrichment for p65 (RelA), p62 (SQSTM1) and HDAC6, which is another aggrephagy associated protein (Fig. 6B-C). Notably, for IL-1β, the volcano plot shows none of the significant partners of COMMD2 held any known NF-kB association (via BioGRID) at 2hrs (Fig. 6D). Moreover, there was a 5- and 80-fold higher recovery of significantly enriched interacting partners under the TIGR4 condition compared to 6B ST90 and IL-1β respectively (Fig. 6B-D; Sup. Table 3). Cross comparing these targets, we discovered that 254 unique proteins were associated with COMMD2 under TIGR4 challenge with 92 of them being also found in our p65 interactome dataset (Fig. 6E; Sup. Table 1). Remarkably overall this unique comparison of the COMMD2 interactome datasets clearly supported the specificity of COMMD2 – p65 (RelA) during TIGR4 challenge.

Finally, analysis using Panther ontology protein class of the significant targets obtained from the interactome underscored a unique TIGR4 enrichment for host metabolite and protein modifying enzymes as well as proteins associated with translation (Fig. 6F). From the analysis we also identified a putative COMMD2 posttranslational modification associated with ubiquitination at Lysine residue 50 (Sup. Table 2), which aligns with our previous endogenous co-immunoprecipitation of COMMD2-p65 (Sup. Fig. 6A). Altogether, these results showed during TIGR4 challenge COMMD2 function was specific with a potential role in host inflammatory response.

### COMMD2 is a selective negative regulator of host cellular immunity

Having previously shown TIGR4 directly represses host transcription and in particularly NF-kB dependent targets ^6,41^, we speculated that COMMD2 was required to alter NF-κB p65 driven host immunity upon stimulation with TIGR4. Thus, inhibiting COMMD2 would lead to increased cytokine production. To our knowledge, no direct chemical inhibitors of COMMD2 exist, so we used RNAi to knockdown COMMD2 and p62 in primary human nasal epithelial cells (Sup. Fig. 9A-C). Whereas both siRNAs to COMMD2 or their mixture resulted in 50-60% reduction, a single siRNA to p62 reduced the target >70% (Sup. Fig. 9C). While knockdown of COMMD2 was not ideal it was consistently greater with the second siRNA, thus we proceeded with bacterial challenge suspecting this minimal reduction in COMMD2, in comparison to scrambled or p62, would be enough to discern roles for COMMD2 and/or p62 upon the analytes within our Luminex panel.

Following 48hrs of either Scrambled (Scr), COMMD2 or p62 RNAi primary human nasal cells were challenged for 4 hrs with either IL-1β, TIGR4 or 6B (± IL-1β), with post-challenge supernatants collected for Luminex (27-plex; COMMD2 Fig. 7A; Scr Sup. Fig. 9D; p62 Sup. Fig. 9E). Strikingly COMMD2 RNAi resulted in selective elevation of multiple targets, even in the uninfected group, with several being significantly higher than their respected scr control (pV reported; Fig. 7A). These analytes - CCL20, CXCL1, CXCL10, IL-1ra, IL-8 and PTX3 - either border or were significantly different with respect to each sample scr control. Interestingly, CCL20, CXCL1, CXCL10, IL-1ra, and IL-8 were also elevated in the p62 RNAi data (Sup. Fig. 9E). Given this surprising result, we determined whether COMMD2 RNAi impacted uninfected, IL-1β, TIGR4 or 6B equally. For this we first compared the absolute difference of COMMD2 RNAi to scr (Fig. 7B). Our results showed that while on average TIGR4 challenge of COMMD2 RNAi cells increased all 27 analytes in comparison to 6B and uninfected cells these elevated levels remained comparable to the IL-1β treated control. However, to assess the degree in which COMMD2 or p62 RNAi affected TIGR4 or 6B we compared each RNAi condition to the scr control (Fig. 7B). This comparison clearly showed knockdown of COMMD2 and p62 during TIGR4 challenge had far more impact on the profiles of the 27 analyte examined than for 6B. Altogether, these data show that: 1) COMMD2, and likely p62, have a role in regulating specific cellular immunity molecules in uninfected cells, and 2) TIGR4 relies on both COMMD2 and p62 to alter secreted host immunity analytes.

Due to this, we explored whether a commonality existed in predicted NF-kB p65 (RelA) binding motifs among these targets transcriptional start site (TSS). Herein, we pulled ± 2kb surrounding the gene TSS (UCSC genome browser) for Transcription factor Affinity Prediction (TRAP) analysis ^47^ and for comparison of the NF-kB motif to the canonical motif (JASPAR). Mapping predicted NF-kB motifs from the COMMD2 targets did reveal several. However, there was no apparent commonality in location or binding affinity of these predicted motif sites in this small dataset (Fig. 7C). However, the NF-kB motif consensus sequence of the small COMMD2 targets, in comparison to the canonical motif using Weblogo ^48^, showed a subtle change with higher probability of G/A (pos. 3) and C/T (pos. 7; Fig. 7D). Interestingly, p62 RNAi also resulted in selective elevation of CCL20, CXCL1, CXCL10, IL-1ra, IL-8, SPI, CXCL2, IL-6, MMP-1 and TNFα in one or more samples (Sup. Fig. 9E). CXCL10, even in uninfected samples, was significantly increased (p value reported) under both COMMD2 and p62 RNAi conditions, potentially suggesting a role of each in its regulation.

Overall, these results are the first evidence that COMMD2 is indeed a selective negative regulator of NF-kB dependent gene transcription. Moreover, analysis of the targeted genes suggests an unknown chromatin level component in the role of COMMD2 – p65 for gene expression.

### COMMD2 nuclear translocation favors invasive pneumococcal strains

Since COMMD2 nuclear localization only occurred during TIGR4 challenge we sought to substantiate whether key TIGR4 virulence determinants were responsible. We first investigated whether Pneumolysin (Ply) or hydrogen peroxide production via Pyruvate oxidase (SpxB) were responsible for COMMD2 nuclear translocation.

Herein, we challenged GFP-COMMD2 cells with the ΔPly or ΔSpxB TIGR4 mutants alongside their parental wildtype strain (Fig. 8A). Comparing ΔSpxB to ΔPly and their parental wildtype showed that a ΔPly mutant does not trigger translocation, whereas the ΔSpxB mutant induced a partial phenotype with ΔSpxB increasing nuclear COMMD2 levels over ΔPly but remaining at 75-80% of wildtype (Fig. 8B). Given this result, we tested if a functional pneumolysin pore was required. For this we used the JWTIGR4 ΔPly W433F pore mutant ^49,50^. This mutant did not induce COMMD2 translocation (Fig. 8B). Altogether we concluded that COMMD2 translocation was dependent on Pneumolysin pore formation in combination with SpxB.

However, the ΔSpxB phenotype was puzzling as the strain retains functional native PLY yet still has reduced COMMD2 levels in comparison to wildtype TIGR4. This disparity suggested that both SpxB and PLY are necessary for COMMD2 nuclear accumulation under wild type conditions. We addressed the synergy between PLY and SpxB by comparing different phenotypes using exogenous purified Pneumolysin toxin alone or in combination with the main byproduct of SpxB, hydrogen peroxide. We first determined the IC_50_ for hydrogen peroxide and purified Pneumolysin toxin individually at 28.30µM and 7.10nM, respectively, however when combined the calculated IC_50_ for hydrogen peroxide dropped to 10µM (Sup. Fig. 10A & B). These results show that the combination of PLY and SpxB impact cellular toxicity more than with each individual component ^51^.

We then tested the impact of exogenous supplementation of 0 – 6.25µM hydrogen peroxide during challenge with either a wildtype, a TIGR4 ΔSpxB, or a TIGR4 ΔSpxBΔPly mutant strain (Sup. Fig. 10D – F). Our results showed increasing concentration of exogenous hydrogen peroxide resulted in TIGR4ΔSpxB phenocopying wildtype except at the highest concentrations, where wildtype induced more nuclear COMMD2. There was no significant increase in nuclear COMMD2 levels for either the TIGR4 ΔSpxBΔPly or JW TIGR4ΔPLY W433F pore mutants against their parental wildtype strains (Sup. Fig. 10E & F). Overall, these data suggest, at least for TIGR4, that PLY and hydrogen peroxide are mutually synergistic in driving nuclear COMMD2 levels.

Capsule is another key bacterial determinant of pneumococcus and drives many virulence mechanisms. We therefore tested whether capsule was required for COMMD2 nuclear translocation. By comparing a wild type (JW TIGR4) to a capsule mutant (JW TIGR4ΔCPS), we observed no difference in their ability to induce nuclear translocation of COMMD2 (Fig. 8E).

We next assessed the association of COMMD2 nuclear localization with isolates favoring either commensal-like/carriage (6B ST138, 11A ST6521 & 19A ST276) or invasive (1 ST304 or 1 ST306 - which harbors a non-hemolytic Pneumolysin toxin) for their capacity to drive COMMD2 nuclear translocation. As expected, our results showed that invasive TIGR4 challenge significantly increased nuclear COMMD2 (pV≤ 0.0001) in comparison to the commensal-like 6B ST90 isolate and uninfected cells (Fig. 8E). Comparing the invasive serotype 1 ST304 and ST306 (non-hemolytic) isolates to TIGR4 revealed no significant differences in their capacity to induce nuclear COMMD2, whereas the commensal-like/carriage 11A ST6521 and 19A ST276 isolates resembled 6B ST90 which did not display COMMD2 localization (Fig. 8E). Interestingly, the two 6B strains, which only differ in their genomic sequence rather than their capsule, were phenotypically different in their nuclear COMMD2 levels. In contrast to 6B ST90 (Global Pneumococcal Sequence Cluster^52^; GPSC 23), 6B ST138 (GPSC 24) not only induced significantly (pV≤ 0.05) more COMMD2 translocation in comparison with the uninfected control but had an elevated level compared to 6B ST90 (pV = 0.07; Fig. 8E). Given this result, we assessed whether the elevated COMMD2 levels with 6B ST138 correlated with in vivo lethality using two challenge doses in an intranasal murine challenge model comparing 6B ST138 to TIGR4 (Fig. 8F & Sup. Fig 10G). At the high dose 6B ST138 induced lethality to a comparable level to that of TIGR4, which stands in contrast to our previously demonstrated findings that 6B ST90 was not lethal in this model even at a log higher dose to that of TIGR4 ^6^. Therefore, we concluded that in a dose dependent manner challenge with 6B ST138 (GPSC 24) has higher pathogenic potential than 6B ST90 (GPSC 23), and this correlates with the differential COMMD2 nuclear signal. Overall, the combination of these data provides strong evidence that COMMD2 translocation is predominantly associated with pneumococcal virulence phenotypes both in vitro and in vivo.

## Discussion

Cellular inflammatory response is a critical component of the host defense to bacteria. Yet, the molecular processes that fine-tune cellular immunity cascades across the range of colonizing to virulent bacteria are poorly understood. We show that an invasive *S. pneumoniae* TIGR4 strain, which causes symptomatic disease in murine models ^6^, drives a different NF-κB p65 activation profile in comparison to a commensal-like 6B ST90 strain and IL-1β – a potent inflammatory cytokine. Our mass spectrometry interactome and post-translational modification analysis, showed these two pneumococcal isolates have diverging p65 interacting partners and phosphorylation status. Analysis of these partners demonstrated that the invasive TIGR4 strain enriched for aggrephagy pathway components and COMMD2, whereas the 6B ST90 strain did not. Through our study of COMMD2 we show that TIGR4 triggers p65 degradation and induces a unique interaction of COMMD2 with p65 and p62. By deeply investigating this process we discovered COMMD2 is a negative regulator of cellular immunity and its nuclear levels associated with invasive favoring pneumococcal isolates. Altogether, we show a bacterial pathogenesis mechanism repressing host cellular immunity through bacterial driven degradation of NF-κB p65 via novel fine-tuning of its interacting partners.

Negative regulation of NF-κB signaling, in contrast to the breadth of knowledge on activatory mechanisms, is poorly documented. This is in part due to the lack of identified targets and mechanisms responsible for attenuating this signaling cascade. Of the known negative regulators, A20 (TNAIP3) and COMMD1 are the better described with the studies using cytokine stimulation. Pointedly neither of these proteins were found in our TIGR4 p65 interactome, which suggests that COMMD2 recruitment during bacterial infection is specific. A20 is primarily a deubiquitinase whose transcription is NF-κB activation dependent. A20 functions in a negative feedback loop to deubiquintinate NEMO, which results in its stabilization with the IKK complex to restore NF-κB sequestration in the cytoplasm. This ultimately terminates the downstream canonical NF-κB signaling cascade of inflammatory response ^18,25,53^. In contrast, COMMD1 transcription is NF-κB independent, and facilitates p65 termination by CRM1 mediated export and translocation of p65 to the proteasome for degradation, while in parallel occupying the formerly p65 bound kappa-binding site at specific gene promoters ^27,29,30,32,34,35^ ^33^. It was put forth that the diversity of potential COMMD and NF-κB assemblies respond to an array of physiological stimuli to fine-tune cellular immunity positioning this family of proteins as powerful selective negative regulators of NF-κB signaling. Interestingly, our post-translational modification analysis of COMMD2 discovered a ubiquitinated peptide associated with a lysine at residue 50, which aligns with our results indicating selective degradation of COMMD2-p65-p62 complex. The COMMD2 negative feedback loop on p65 that we uncover here may represent, even in a cellular state without bacterial infection, a precise mechanism to terminate or shift a given p65 dependent transcription repertoire.

Given the vast array of post-translational modifications on p65 and other NF-κB subunits across differential stimulations has given rise to the “NF-κB barcode hypothesis”, which suggests that distinct patterns are linked to how inflammatory gene transcription occurs ^54,55^. Supporting this is our exploratory mass spectrometry of p65 phosphorylation, which identified serine 45 (S45) as the only enriched phosphorylated mark during TIGR4 challenge. This mark has previously been shown to negatively regulate p65, although the mechanism is unknown ^56^. Lanucara et al., showed that a phosphomimetic mutant of S45 prevented IL-6 transcription and p65 binding to the IL-6 promoter after 3 hours of TNFα stimulation ^56^. It remains to be evaluated how this modification affects COMMD2-p62 degradation of p65 or upon repressing p65 host transcription. Notably our previous studies showed TIGR4 challenge induces de-phosphorylation of histone H3 serine 10, which is an essential histone modification associated with inflammatory gene transcription ^41,57^. Understanding how this event relates to p65 turnover and COMMD2 may shed light on essential chromatin remodeling dynamics needed to precisely regulate NF-κB p65 DNA binding, extraction and transcription termination not only to pneumococci but also under inflammatory stimuli. Excitingly, the commensal-like 6B ST90 does not induce phosphorylation of this S45 residue. Instead, this strain leads to phosphorylation on S203 in addition to S536 and S276. These marks may be critical for the upregulation of the histone demethylase KDM6B, which we previously showed was required for bacterial containment in the upper respiratory tract ^6^. Future investigations here may be the best positioned to decipher outstanding questions on how bacteria engage the non-canonical NF-κB pathway and how KDM6B is selectively recruited to gene targets during bacterial infection. Whether differential phosphorylation of p65 is the determining factor in the host response to different strains of pneumococcus remains to be determined. It is tempting to speculate that posttranslational modifications of p65 may represent markers of either host response to commensal or to invasive bacteria.

To date our work is the first to show that COMMD2 mediates p65 turnover through p62 and aggrephagy, a selective form of autophagy, while simultaneously repressing host cellular immunity. TIGR4, along with other pneumococcal isolates associated with higher virulence potential, triggered COMMD2 nuclear accumulation. These findings suggest that nuclear localization of COMMD2 could be a marker of virulence, distinguishing the host response from commensal-like isolates. We show that COMMD2 nuclear translocation was independent of the TIGR4 capsule, but dependent on a tandem function of Pneumolysin (Ply) and Pyruvate oxidase (SpxB), as a ΔPly mutant shows no COMMD2 nuclear accumulation, but a ΔSpxB mutant induced partial COMMD2 translocation with a distinct phenotype. Our efforts titrating hydrogen peroxide, the product of *SpxB*, with the TIGR4 ΔSpxBΔPly or JW TIGR4ΔPLY W433F pore mutants revealed at least for TIGR4, PLY and hydrogen peroxide are mutually synergistic in driving nuclear COMMD2 levels and strongly support that Pneumolysin may be “masking” other bacterial effectors. Given the documented influence of Pneumolysin toxin, like other CDC toxins, on host plasma membrane, the known differences in Pneumolysin alleles ^58^, and the known role of COMMD proteins, along with CCDC22 and CCDC93 (the CCC complex), in phosphatidylinositol-3-phosphate (PI3P) equilibrium within endosomes ^44^ raises the possibility that combinations of Pneumolysin toxin alleles with hydrogen peroxide may alter host membrane compositions. Deeper comparative studies between pneumococcal isolates and virulence associated bacterial protein mutants may likely identify new bacterial proteins that alter host response. Moreover, our comparison of two serotype 6B isolates varying in sequence type and Global Pneumococcal Sequence Cluster resulted in different COMMD2 nuclear amounts which correlated with increased 6B ST138 (GPSC 24) dose dependent lethality during intranasal murine challenge. Thus, we propose invasive favoring pneumococcal isolates, as opposed to commensal-like ones, shape host cellular immunity, and potentially disease onset and lethality, through potent shifts in host transcription factor partners.

Tuning cellular immunity is fundamental for processes of airway epithelial cells exposed to pneumococcus ^7,59^. The pro-inflammatory cytokines, TNF-α and IL-1β, are major cytokines necessary for neutrophil recruitment and are found in bronchoalveolar lavage fluid of animals challenged with pneumococcal isolates ^7,59,60^. However, a study showed that isolated murine lung epithelial cells exposed to serotypes 19 and 3 failed to induce p65 (RelA) nuclear translocation in comparison to TNFα and IL-1β cytokine treatment alone ^7^. Indeed during respiratory infection with other microbes ^61,62^, a balance is needed between pro-inflammatory responses and negative regulation to ensure minimal tissue damage from the influx of neutrophils into the airway tissues ^60^. Airway epithelial cells play a crucial role in both situations by regulating neutrophil recruitment and promoting epithelial repair pathways leading to tissue resilience and resolution of inflammation ^59,60,63,64^. With invasive pneumococci actively antagonizing the ability of airway epithelial cells to mount a cellular immune response, we hypothesize an amplifying and runaway inflammatory cascade is created in latter stages of infection where neutrophil influx is detrimental ^59,64,65^. This could lead to exacerbated and severe pneumonia with excessive tissue damage, allowing pneumococcus to transmigrate to deeper tissues. We put forth that COMMD2, or combinations of COMMD proteins are modulators of bacterial driven inflammatory processes and may represent a therapeutic target to circumvent collateral tissue damage.

In conclusion, our study shows a new regulatory role for COMMD2 in restraining p65 through aggrephagy mediated turnover triggered by bacterial interaction. We reveal this process to be specific to invasive TIGR4 pneumococcal challenge and partially depend on pneumolysin. Further studies characterizing both the p65 and COMMD2 interactome under bacterial challenge with isolates representing divergent pneumococcal host interaction may identify new processes exploited at the microbe-host interface to regulate cellular immunity.

## Materials and methods

### Bacteria strains, growth, and enumeration

Pneumococcal isolates of 6B ST90 CC156 lineage F (ST90), 6B ST138, 11A ST6521, 19A ST276, 1 ST304 or 1 ST306, and TIGR4 were obtained from either the Centre National de Référence des Pneumocoques (Emmanuelle Varon; Paris, France) or (Thomas Kohler, Universität Greifswald) respectively. JW TIGR4 and ΔW433F pore mutants were a gift from Dr. Jeffrey N. Weiser ^49^. The TIGR4 ΔSpxB and TIGR4 ΔSpxBΔPly mutants were generated previously ^41^. Experimental starters were made from master glycerol stocks struck on 5% Columbia blood agar plates (Biomerieux Ref# 43041) and grown overnight at 37°C with 5% CO_2_ prior to outgrowth in Todd-Hewitt (BD) or Brain heart infusion (BHI) broth as previously described ^6^. Inocula were prepared from frozen experimental stocks grown for 3 – 4 hrs to midlog phase in BHI at 37°C with 5% CO_2_ in closed falcon tubes. Bacterial cultures were pelleted at 1,500xg for 10 mins at room temperature (RT), washed in DPBS, and concentrated in 1mL DPBS prior to dilution at desired CFU/mL using 0.6 OD_600_ /mL conversion factors in desired cell culture media ^6^. Bacterial counts were determined by serial dilution plating on 5% Columbia blood agar plates and grown overnight at 37°C with 5% CO_2_.

### Cell culture conditions and in vitro challenge

A549 human epithelial cells (ATCC ref# CCL-185) and A549 stable cell lines were maintained in F12K media (Gibco) supplemented with 1x GlutaMax (Gibco) and 10% heat inactivated fetal calf serum (FCS) at 37°C with 5% CO_2_. Primary human nasal epithelial cells (HNEpC; PromoCell ref# C-12620; lot# 475Z023) were cultured and maintained in PneumaCult™-Ex Plus Medium (STEMCELL Technologies cat# 16237380). All cell lines were discarded after passage 4. For challenge studies cells were plated in tissue culture treated plates at 2×10^5^ cells (6well; for 72 hrs), 5×10^4^ cells (24well; for 48 hrs), or 1×10^4^ cells (96well; for 48 hrs) ^66^. Bacterial inocula (Multiplicity of infection (MOI) 20) were diluted in cell culture media, added to cells, and bacterial-epithelial cell contact synchronized by centrifugation at 200xg for 10 mins at RT. Plates were moved to 37°C with 5% CO_2_ for 2 hrs and processed as desired for experiment termination. For inhibitor studies, cell culture media was aspirated, and replaced with filter sterilized culture media containing either Bafilomycin A1 400 nM final concentration (Sigma ref# SML1661) or Leptomycin B 10 nM final concentration (Sigma ref# L2913) for 3 hrs prior to bacterial addition. Human IL-1β (Enzo Life Sciences ref# ALX-522-056) was used at 10 ng/mL final concentration in cell culture media.

### RNA isolation and RT-qPCR

Total RNA isolated and extracted using TRIzol (Life technologies ref#15596-026) method as per manufacturer’s recommendations. Recovered RNA (5 µg) was converted to cDNA with Super Script IV as per manufacturer’s instructions, diluted to 20 ng/µL in molecular grade water and 1 µL used for Sybr Green reactions as per manufacturer’s instructions on a BioRad CFX384 (BioRad). Relative expression was calculated by ΔΔCt method to *GapDH* ^67^ with RT-PCR primers in Sup. Table 3 ^6^.

### ChIP and ChIP-qPCR

Detailed ChIP buffer components and procedure were completed as previously reported ^66^. Briefly, 8×10^6^ A549 cells were cross-linked with 1% formaldehyde at room temperature and quenched with 130 mM glycine. Chromatin was generated from the collected cell pellets by lysis and sonication in chromatin shearing buffer to a size of 200-900bp. ChIP grade antibody to p65 (L8F6) (CST ref #6956) was used at manufacturer’s recommended concentrations and bound to DiaMag beads (diagenode ref # C03010021-150) overnight with gentle rotation. Quantified chromatin was diluted to 10 µg per immunoprecipitation and added to antibody bound DiaMag beads overnight with gentle rotation and 8% of input reserved. Beads were washed as previously described ^66^, and DNA purified using phenol-chloroform extraction followed by isopropanol precipitation. Recovered DNA suspended in molecular grade water was used for Sybr Green reactions (1 µL) on a BioRad CFX384 (BioRad). ChIP-qPCR primers (50-150 bp; 60 °C max melt temperature) were designed to span the NF-κB sites of interest within the promoters of *CFS2* or *PTGS2* ^68^. % recovery was calculated as 2 raised to the adjusted input Ct minus IP Ct multiplied by 100. ChIP qPCR primers listed in Sup. Table 3.

### Plasmids, molecular cloning and stable cell line generation

All plasmids and primers are listed in Sup. Table 3. Routine cloning was carried out by in vivo assembly ^69,70^. Briefly, primers were designed with a 15-20 bp overlap to amplify nucleic acid targets using Phusion Plus polymerase (Thermo ref# F630S). Correct sized bands were excised and nucleic acid extracted by “Freeze and squeeze” ^71,72^. Herein, 0.7% - 1% agarose gel fragments were frozen for 5 mins on dry ice and centrifuged for 15 mins at >21,000 xg with the supernatant collected – the process was completed two additional times. Collected supernatant containing nucleic acid was then purified using phenol-chloroform extraction followed by isopropanol precipitation and suspension in molecular grade water. Collected nucleic acid was quantified spectrophotometrically using a NanoDrop and mixed at 3:2 (vector: insert) in 10 µl and added to chemically competent E. coli MC1061 or DH5α for transformation. After 1 hr incubation on ice bacteria outgrowth was done for 1 hr in Luria-Bertani (BD) prior to selection on LB agar containing Ampicillin antibiotic (100 µg/mL). All plasmids were isolated with the QIAprep Spin Miniprep Kit (Qiagen ref# 27106) and eluted in molecular grade water (endotoxin free) as per manufacturer’s instructions. A549 stable cell lines were generated using the transposon-based sleeping beauty system ^73,74^. A549 cells were plated in tissue culture treated plates at 2×10^5^ cells (6well) one day prior to transfection with 2 µg plasmid DNA + 150 ng SB100 transposase DNA. After transfection, cells were selected with 1 mg/mL Geneticin (Thermo ref# 10131035) for 7 days, with media exchanged on days 1, 3, 5 & 7. Cells were collected with Trypsin 0.25% EDTA (Thermo ref# 25200056) and two-way serial diluted in a 96 well tissue culture plate for another 7 – 14 days with 1 mg/mL Geneticin to select colonies that were phenotypically bright were mixed and expanded prior to FACS sorting to ensure purity, and uniform expression.

### Immunoblots and quantification

Whole cell lysates were obtained by RIPA lysis (10 mM Tris HCL pH 7.5, 150 mM EDTA, 0.1% SDS, 1% Triton X-100 & 1% Deoxycholate) supplemented with inhibitor cocktail (1X PhosSTOP, 10 mM sodium butyrate, 0.2 mM PMSF). Samples combined with 5x with Laemmli buffer ^75^, sonicated for 5 mins in a ultrasonic water bath, boiled at 98°C (dry bath) for 10 mins and frozen at −20°C. Whole cell lysates were ran on 4 – 20% pre-cast polyacrylamide SDS PAGE gels (BioRad), transferred to PVDF membrane (BioRad TransBlot) and blocked 1 hr in 5% BSA TBST at room temperature. Membranes were probed overnight at 4°C in 5% BSA TBST with primary antibody to p65 (CST ref #6956 or CST ref# 8242), p65 phosphorylation at serine 536 (CST ref# 3033), p65 phosphorylation at serine 276 (abcam ref# ab183559), SQSTM1 / p62 antibody [EPR4844] (abcam ref# ab109012) or actin AC-15 monoclonal (Sigma ref# A5441) as per manufacturer’s recommendations. COMMD2 immunoblots were blocked for 1 hr at room temperature in 3% milk TBST and probed overnight at 4°C with the primary antibody (Sigma ref# HPA044190) suspended in 3% milk TBST. Appropriate secondary-HRP conjugated antibodies in 5% Milk TBST were incubated for 1 hr at room temperature prior to development with clarity ECL (BioRad) developing reagents with a ChemiDoc Touch (BioRad). Band intensity was quantified by Image Lab (BioRad) with linear intensity values log_10_ transformed and normalized to actin prior to any additional ratio metric comparisons.

### siRNA and transfection

Primary human nasal cells were seeded at 2×10^4^ cells per well in a 96well plate the day before RNAi transfection. For RNAi 1 µl of 2 µM Silencer Select siRNA stock for either COMMD2 (Ambion ID# s27490; or Ambion ID# s27488) or SQSTM1 (p62; Ambion ID# s16961) was diluted with 9 µl Opti-MEM (Thermo cat# 31985062) and mixed with 10 µl of 0.003% (vol/vol) Lipofectamine RNAiMAX (Thermo cat# 13778030)/ Opti-MEM. For COMMD2 siRNA mix, a final 2 µM concentration of siRNA was respected. Complexes were formed during 15 min incubation at room temperature prior to 20 µl of mixture added to previously seeded cells. 96well plate was incubated for an additional 72 hrs 37°C with 5% CO_2_ prior to challenge studies.

### Cell fractionation

Fractionation was performed as previously described as previously described ^66^. Faction lysates were combined with 5x with Laemmli buffer ^75^, sonicated for 5 mins in a ultrasonic water bath, boiled at 98°C (dry bath) for 10 mins and frozen at −20°C. Samples were ran on either 10% (for GFP-COMMD2) or 12% (for fraction quality controls) polyacrylamide SDS PAGE gels (BioRad), transferred to PVDF membrane (BioRad TransBlot), blocked 1 hr in 5% BSA TBST at room temperature. Primary antibody in 5% BSA TBST to GFP (abcam ref# ab290), GapDH (abcam ref# ab8245), or histone H4 (abcam ref# ab177840) was completed overnight at 4°C. After 3x 10 min washes in TBST appropriate secondary-HRP conjugated antibodies in 5% Milk TBST were incubated for 1 hr at room temperature and developed with a ChemiDoc Touch (BioRad) as described above.

### Immunofluorescence microscopy and Cellprofiler analysis

For microscopy cells were seeded either on acid washed and UV treated coverslips (24well) or directly in 96well plates as described above. Two hours post-challenge media was aspirated, cells washed in DPBS, and fixed with 2.5% PFA for 10 mins at RT. Fixed cells were blocked and permeabilized overnight in 5% BSA 0.5% Tween20 at 4°C. Primary antibody to p65 (CST ref #6956 or CST ref# 8242), or p62 (SQSTM1; abcam ref# ab109012) were diluted at 1:1,000 in 5% BSA 0.5% Tween20, for time course of p65 phosphorylation at serine 536 cells were stained with MA5-15160 (Thermo; 1:1000), and incubated overnight at 4°C. Cells were washed 3x 10 mins at RT in PBS + 0.1% Tween20 prior to 1 hr incubation at 1:1,000 dilution of either Alexa Fluor 594 or Alexa Fluor 647 secondary antibody. Nuclei were stained with 10 ng/mL final concentration of Hoechst 33342 or Draq5 for 15 mins. Coverslips were rinsed in PBS and molecular grade water prior to mounting with Fluoromount-G Mounting Medium (INTERCHIM). Confocal microscopy images were acquired on a Nikon TiE inverted microscope with an integrated Perfect Focus System (TI-ND6-PFS Perfect Focus Unit) and a Yokogawa Confocal Spinning disk Unit (CSU-W1). Nine images per well were acquired using a 20X air objective (NA 0.75) at a step-size of 0.9µm in z-plane. Deconvoluted epifluorescent images were acquired on a Cytation 5 (BioTek) using a 20X air objective (NA 0.75) with a grid of 3 x 3 (9 images en total).

Images were processed for background using Fiji ^76^, and segmented using Cell Profiler ^77–79^. Briefly, image analysis consisted of sequential modules to ‘IdentifyPrimaryObjects’ based on channel signal for nuclei (Hoechst 33342 or Draq5 stain), p65 (Alexa594), or p62 (Alexa594). This was followed by ‘IdentifySecondaryObjects’ for the GFP-COMMD2 signal via propagation of identified nuclei. Objects were related to each other to maintain cohesion between identified nuclei, cell and cellular contents (p65 or p62). For puncta, the additional module, ‘EnhanceorSupressFeatures’ with ‘Speckles’, was used. This used a global threshold strategy with Otsu threshold method and a 2% minimum boundary to identify puncta contained within the segmented nuclei.

### Immunoprecipitation

Cells were lysed in 250 μL of RIPA lysis (10 mM Tris HCL pH 7.5, 150 mM EDTA, 0.1% SDS, 1% Triton X-100 & 1% Deoxycholate) supplemented with a protease mixture inhibitor (Roche Complete, EDTA free). Lysates were either immunoprecipitated using GFP-trap agarose beads (ChromoTek ref# gta-10) or with DynaBeads Protein G (Thermo ref# 10004D). For GFP-p65 and GFP-COMMD2 the samples were immunoprecipitated as per manufacturer’s instructions with the elution was recovered in either 5x with Laemmli buffer ^75^ and boiled at 98°C (dry bath) for 10 mins, or left in Trypsin digest buffer (see LC-MS/MS Mass-spectrometry and analysis). All samples were frozen at - 20°C. For endogenous samples the lysates were incubated on a rotating wheel at 4 °C for 20 min before adding 1 mL of dilution buffer (150 mM NaCl and 50 mM Tris-HCl pH 7.5 supplemented with Protease mixture inhibitor) to reduce the detergent final concentration below 0.1%. The lysates were then centrifuged at 10,000 × g for 10 min. For p65 IP the lysates were then incubated with 2 μg of antibody CST ref #6956) at 4 °C overnight before adding 50 μL of Dynabeads for 1 hr. The beads were then washed three times in 0.01%Tween20 PBS before adding 20 μL of Laemmli buffer supplemented with 2% β-mercaptoethanol and boiled for 5 min.

### Exogenous Hydrogen peroxide and Pneumolysin toxin challenges

For viability studies titrations of hydrogen peroxide (9 M stock, Thermo ref# 411881000; 0 µM – 240 µM) and Pneumolysin toxin (50µM stock; gift Dr. T. Mitchell; final 0 nM – 50 nM), concentrations were serial diluted into cell culture media. After 2 hrs of incubation samples were PFA fixed and nuclei stained with Draq5 (Thermo ref# 62252) to determine viability. Bacterial challenge experimental starters of TIGR4 ΔSpxB, TIGR4 ΔSpxBΔPly, JW TIGR4 and ΔW433F were grown and diluted in cell culture media as in “Bacterial strains, growth and enumeration”. These inocula were combined with cell culture media containing indicated hydrogen peroxide concentrations (0 µM – 6.25 µM) and added to cells seeded in 96 well plates. After 2 hrs cells were PFA fixed and COMMD2 nuclear levels determined as in “Immunofluorescence microscopy and Cellprofiler analysis”.

### Cytokine quantification by Luminex

Supernatants we collected from primary nasal epithelial cells seeded in a 96well plate as previously described. For this 200 µl of supernatant per well was collected 2 hrs post-challenge with IL-1β or bacteria, centrifuged in a 96well U-bottom plate for 10 min at 1,500 x g to pellet bacteria and cellular debris. 120 µl of cleared supernatant was collected into a 96well flat bottom plate and frozen at −20°C until use. Secretion in the supernatant of 29 proteins (Human Magnetic Luminex Assays, R&D Systems) was measured following manufacturer’s instructions using Luminex xMAP technology. Briefly, samples and standards were incubated with capture beads for 2 hours at RT. After washing, detection antibody mix was added and incubated for 1 hour at RT. After washing, streptavidin-PE was added and incubated for 30 minutes at RT. Samples were acquired on a Bioplex 200 (Biorad). Analysis was performed using the Bioplex Manager Software. Concentration of each analyte was calculated using a 5-PL regression curve.

### LC-MS/MS Mass-spectrometry and analysis

For label-free quantitative proteomic analysis of GFP-p65 and GFP-COMMD2 the respected A549 cell lines were plated in 6well tissue culture plates, and challenged with bacteria for 2hrs as described above. One plate (∼5×10^7^ cells) per condition was harvested using RIPA lysis and immunoperciptated with GFP-trap agarose beads (ChromoTek ref# gta-10) as per manufacturer’s instructions. Three or four independent biological replicates were prepared and analyzed for each condition. Prior to on-bead Trypsin digestion, the samples were washed 3x in trypsin digest buffer (20 mM Tris.HCl pH 8.0, 2 mM CaCl_2_).

For pull down of GFP-p65 on bead digestion was performed strictly as described by Chromotek. Briefly, beads were suspended in digestion buffer (Tris 50 mM pH 7.5, urea 2 M, 1 mM DTT and 5 µg.µl of trypsin (Promega)) for 3 min at 30°C. Supernatants were transfer to new vials and beads were washed twice using (Tris 50 mM pH 7.5, urea 2 M and iodoacetamide 5 mM). All washes were pulled and incubated at 32°C for overnight digestion in the dark. Peptides were purified using C18 stage tips protocol ^80^.

LC-MS/MS analysis of digested peptides was performed on an Orbitrap Q Exactive Plus mass spectrometer (Thermo Fisher Scientific, Bremen) coupled to an EASY-nLC 1200 (Thermo Fisher Scientific). A home-made column was used for peptide separation (C_18_ 30 cm capillary column picotip silica emitter tip (75 μm diameter filled with 1.9 μm Reprosil-Pur Basic C_18_-HD resin, (Dr. Maisch GmbH, Ammerbuch-Entringen, Germany)). It was equilibrated and peptide were loaded in solvent A (0.1 % FA) at 900 bars. Peptides were separated at 250 nl.min^-1^. Peptides were eluted using a gradient of solvent B (80% ACN, 0.1 % FA) from 3% to 31% in 45 min, 31% to 60% in 17 min, 60% to 90% in 5 min (total length of the chromatographic run was 82 min including high ACN level step and column regeneration). Mass spectra were acquired in data-dependent acquisition mode with the XCalibur 2.2 software (Thermo Fisher Scientific, Bremen) with automatic switching between MS and MS/MS scans using a top 12 method. MS spectra were acquired at a resolution of 70000 (at *m/z* 400) with a target value of 3 × 10^6^ ions. The scan range was limited from 300 to 1700 *m/z*. Peptide fragmentation was performed using higher-energy collision dissociation (HCD) with the energy set at 27 NCE. Intensity threshold for ions selection was set at 1 × 10^6^ ions with charge exclusion of z = 1 and z > 7. The MS/MS spectra were acquired at a resolution of 17500 (at *m/z* 400). Isolation window was set at 1.6 Th. Dynamic exclusion was employed within 30 s. Recorded spectra were searched with MaxQuant (version 1.5.3.8) using the Andromeda search engine^81^ against a human database (74368 entries, downloaded from Uniprot the 27^th^ of September 2019), a Streptococcus pneumoniae R6 database (2031 entries, downloaded from Uniprot the 1^st^ of January 2020) and a Streptococcus pneumoniae serotype 4 database (2115 entries, downloaded from Uniprot 1^st^ of January 2020).

The following search parameters were applied: carbamidomethylation of cysteines was set as a fixed modification, oxidation of methionine and protein N-terminal acetylation were set as variable modifications. The mass tolerances in MS and MS/MS were set to 5 ppm and 20 ppm respectively. Maximum peptide charge was set to 7 and 5 amino acids were required as minimum peptide length. At least 2 peptides (including 1 unique peptides) were asked to report a protein identification. A false discovery rate of 1% was set up for both protein and peptide levels. iBAQ value was calculated. The match between runs features was allowed for biological replicate only.

For pull down of GFP-COMMD2, washed beads were re-suspended in 150 µl digestion buffer (20 mM Tris.HCl pH 8.0, 2 mM CaCl_2_) and incubated for 4 hours with 1 µg trypsin (Promega) at 37°C and shaking at 1200 rpm. Beads were removed, another 1 µg of trypsin was added and proteins were further digested overnight at 37°C, shaking at 750 rpm. Peptides were acidified with trifluoroacetic acid (TFA) to lower the pH below 3 and desalted on reversed phase (RP) C18 OMIX tips (Agilent). The tips were first washed 3 times with 100 µl pre-wash buffer (0.1% TFA in water/acetonitrile (ACN, 20:80, v/v)) and pre-equilibrated 5 times with 100 µl of wash buffer (0.1% TFA in water) before the sample was loaded on the tip. After peptide binding, the tip was washed 3 times with 100 µl of wash buffer and peptides were eluted twice with 100 µl elution buffer (0.1% TFA in water/ACN (40:60, v/v)). The combined elutions were transferred to HPLC inserts and dried in a vacuum concentrator. The peptides were re-dissolved in 20 µl loading solvent A (0.1% TFA in water/ACN (98:2, v/v)) of which 2 µl was injected for LC-MS/MS analysis on an Ultimate 3000 RSLCnano system in-line connected to a Q Exactive HF mass spectrometer (Thermo). Trapping was performed at 10 μl/min for 4 min in loading solvent A on a 20 mm trapping column (made in-house, 100 μm internal diameter (I.D.), 5 μm beads, C18 Reprosil-HD, Dr. Maisch, Germany). The peptides were separated on an in-house produced column (75 µm x 250 mm), equipped with a laser pulled electrospray tip using a P-2000 Laser Based Micropipette Puller (Sutter Instruments), packed in-house with ReproSil-Pur basic 1.9 µm silica particles (Dr. Maisch). The column was kept at a constant temperature of 50°C. Peptides eluted using a non-linear gradient reaching 33% MS solvent B (0.1% FA in water/acetonitrile (2:8, v/v)) in 60 min, 55% MS solvent B in 75 min and 70% MS solvent B after 90 min at a constant flow rate of 300 nl/min. This was followed by a 5-minutes wash at 70% MS solvent B and re-equilibration with MS solvent A (0.1% FA in water). The mass spectrometer was operated in data-dependent mode, automatically switching between MS and MS/MS acquisition for the 12 most abundant ion peaks per MS spectrum. Full-scan MS spectra (375-1500 m/z) were acquired at a resolution of 60,000 in the Orbitrap analyzer after accumulation to a target value of 3,000,000. The 12 most intense ions above a threshold value of 13,000 were isolated (isolation window of 1.5 m/z) for fragmentation at a normalized collision energy of 30% after filling the trap at a target value of 100,000 for maximum 80 ms. MS/MS spectra (145-2,000 m/z) were acquired at a resolution of 15,000 in the Orbitrap analyzer. The polydimethylcyclosiloxane background ion at 445.120028 Da was used for internal calibration (lock mass) and QCloud ^82^ has been used to control instrument longitudinal performance during the project. Recorded spectra were searched with MaxQuant (version 2.0.3.0) using the Andromeda search engine with default search settings including a false discovery rate set at 1% on PSM, peptide and protein level.

Spectra were searched against the *Streptococcus pneumonia setotype 4* (TIGR4) (Taxid: 170187) protein sequences in the Uniprot database (database release version of 04_2022, UP000000585), containing 2,109 sequences (www.uniprot.org), the *Streptococcus pneumoniae* (Taxid: 1313) protein sequences in the Uniprot database (database release version of 04_2022, UP000046310), containing 2,115 sequences (www.uniprot.org) and the human protein sequences in the Swiss-Prot database (database release version of 2022_01, reference proteome: UP000005640), containing 20,588 sequences (www.uniprot.org). The mass tolerance for precursor and fragment ions was set to 4.5 and 20 ppm, respectively, during the main search. Enzyme specificity was set as C-terminal to arginine and lysine, also allowing cleavage at proline bonds with a maximum of three missed cleavages. Variable modifications were set to oxidation of methionine residues, acetylation of protein N-termini and lysine residues, phosphorylation of serine, threonine and tyrosine and ubiquitination of lysine residues. Matching between runs was enabled with a matching time window of 0.7 minute and an alignment time window of 20 minutes. Only proteins with at least one unique or razor peptide were retained leading to the identification of 2,328 proteins. Proteins were quantified by the MaxLFQ algorithm integrated in the MaxQuant software. A minimum ratio count of two unique or razor peptides was required for quantification.

### Data analysis for quantitative proteomics

Quantitative analysis was based on pairwise comparison of protein intensities. Values were log-transformed (log2). Reverse hits and potential contaminant were removed from the analysis. Proteins with at least 2 peptides were kept for further statistics after removing shared proteins from the uninfected GFP alone control. Intensity values were normalized by median centering within conditions (normalized function of the R package DAPAR ^83^). Remaining proteins without any iBAQ value in one of both conditions have been considered as proteins quantitatively present in a condition and absent in the other. They have therefore been set aside and considered as differentially abundant proteins. Next, missing values were imputed using the impute. MLE function of the R package imp4p (https://rdrr.io/cran/imp4p/man/imp4p-package.html). Statistical testing was conducted using a limma t-test thanks to the R package limma ^84^. An adaptive Benjamini-Hochberg procedure was applied on the resulting p-values thanks to the function adjust.p of R package cp4p^85^ using the robust method described in (^86^) to estimate the proportion of true null hypotheses among the set of statistical tests. The proteins associated to an adjusted p-value inferior to a FDR level of 1% have been considered as significantly differentially abundant proteins.

### Statistical analysis

All experiments, unless otherwise noted, were biologically repeated 3–5 times and the statistical test is reported in the figure legend. Data normality was tested by Shapiro-Wilk test, and appropriate parametric or non-parametric tests performed depending on result. P values calculated using GraphPad Prism software and the exact values are in source data. Microscopy data obtained from analysis of 3 – 5 image fields per biological replicate after being automatically acquired by the microscope software to ensure unbiased sampling with the total number of analyzed cells or nuclei noted in the figure legend.

## Data Availability

All data in the present study is available upon request from the corresponding authors. The mass spectrometry proteomics data of GFP-p65 and GFP-COMMD2 have been deposited to the ProteomeXchange Consortium via the PRIDE partner repository with the dataset identifiers PXD032970 (p65) and PXD043886 (COMMD2).

## Code Availability

No custom code or software was used in the manuscript.

## Acknowledgements

We would like to thank Emmanuelle Varon and Thomas Kohler for their generous gifts of *S. pneumonie* strains. We are appreciative of Pierre-Henri Commere and the Institut Pasteur, Flow Cytometry Platform (Paris, France) for sorting of the COMMD2 stable cell line. Finally, we would like to thank Daniel Hamaoui for his help processing blots during COVID-19 related work personnel restrictions. Michael G. Connor is supported by a Springboard to Independence grant (AirwayStasis) from the French Government’s Investissement d’Avenir program, the Laboratoire d’Excellence ‘‘Integrative Biology of Emerging Infectious Diseases” (ANR-10-LABX-62-IBEID). Melanie A. Hamon received support from the French government through the National Research Agency (ANR) as part of the France 2030 program referenced “ANR-23-CHBS-0001” (ChromaBac). Work in the laboratory Chromatin and infection unit (headed by Melanie A. Hamon) is supported by the Institut Pasteur, the Fondation pour la Recherche Médicale (FRM-EQU202003010152), the Fondation iXCore-iXLife and the Pasteur-Weizmann research fund. Sebastian Baumgarten is supported by the Institut Pasteur and the European Commission (ERC-STG “PlasmoEpiRNA”). The views expressed in this article are those of the authors and not necessarily those of the NIHR, or the Department of Health and Social Care. Jost Enninga and Lisa Sanchez, members of Dynamics of host-pathogen interactions unit (Institut Pasteur), are supported by the European Commission (ERC-CoG-Endosubvert), the ANR-HBPsensing, and are members of the IBEID and Milieu Interieur LabExes. Filipe Carvalho was supported by a postdoctoral grant through the “Investissement d’Avenir” as part of a French “Laboratoire d’Excellence” (LabEx) research program: Integrative Biology of Emerging Infectious Diseases (ANR-10-LBX-62 IBEID). This work was also supported by EPIC-XS, project number 823839, funded by the Horizon 2020 program of the European Union.

## Author contributions

Conceived and designed all experiments: MGC and MAH. Performed and analyzed data for all experiments: MGC with specific contributions from LS (confocal microscopy imaging); FC, MGE, & TC (p65 mass spectrometry data, repeats for GFP & endogenous immunoprecipitations validations, analysis…); FI, SD and SB (COMMD2 mass spectrometry data, and analysis…), and CC (western blot and cell culture). MGC and TMNC conducted murine challenge models. MGC and MAH edited and reviewed the manuscript. MC and MAH supervised the research and secured funding. All authors approved the final manuscript.

---

**Sup. Figure 1:**
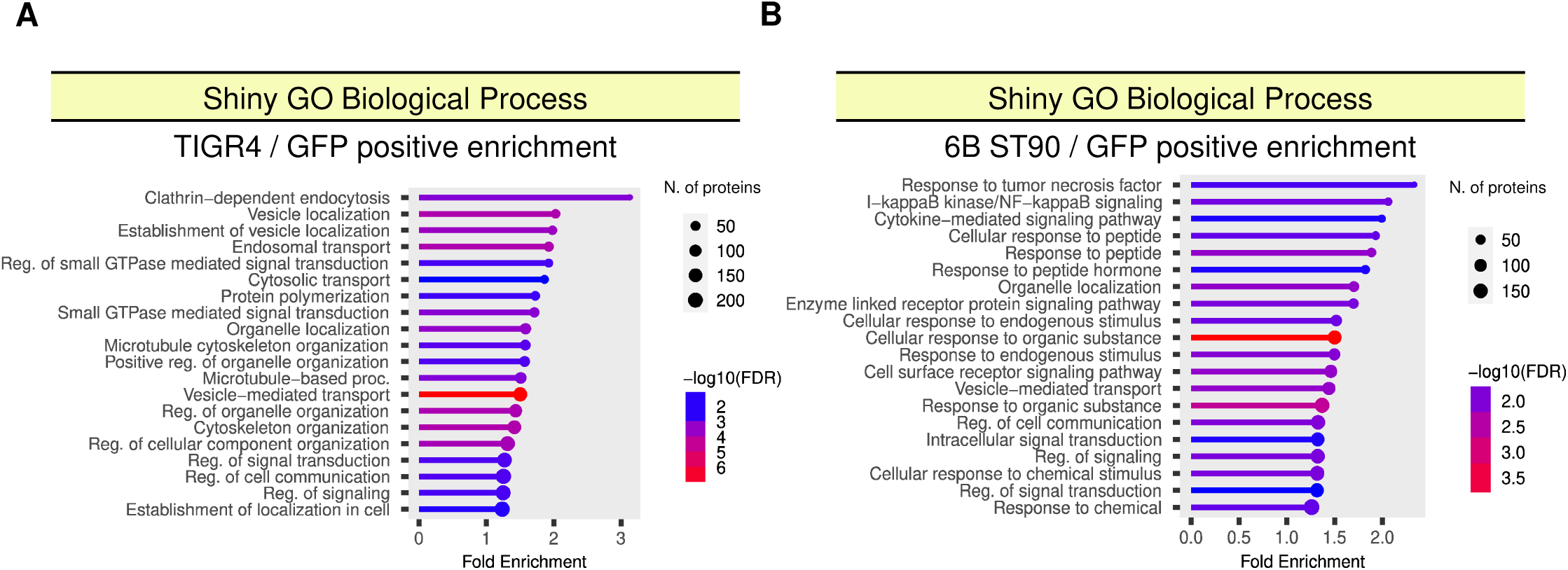
Individual GO biological process enrichment for TIGR4 and 6B ST90 p65 interactomes. A-B) TIGR4 or 6B ST90 enrichment over GFP. Lollipop graph indicating FDR by color scale and number of proteins by lollipop size.

**Sup. Figure 2:**
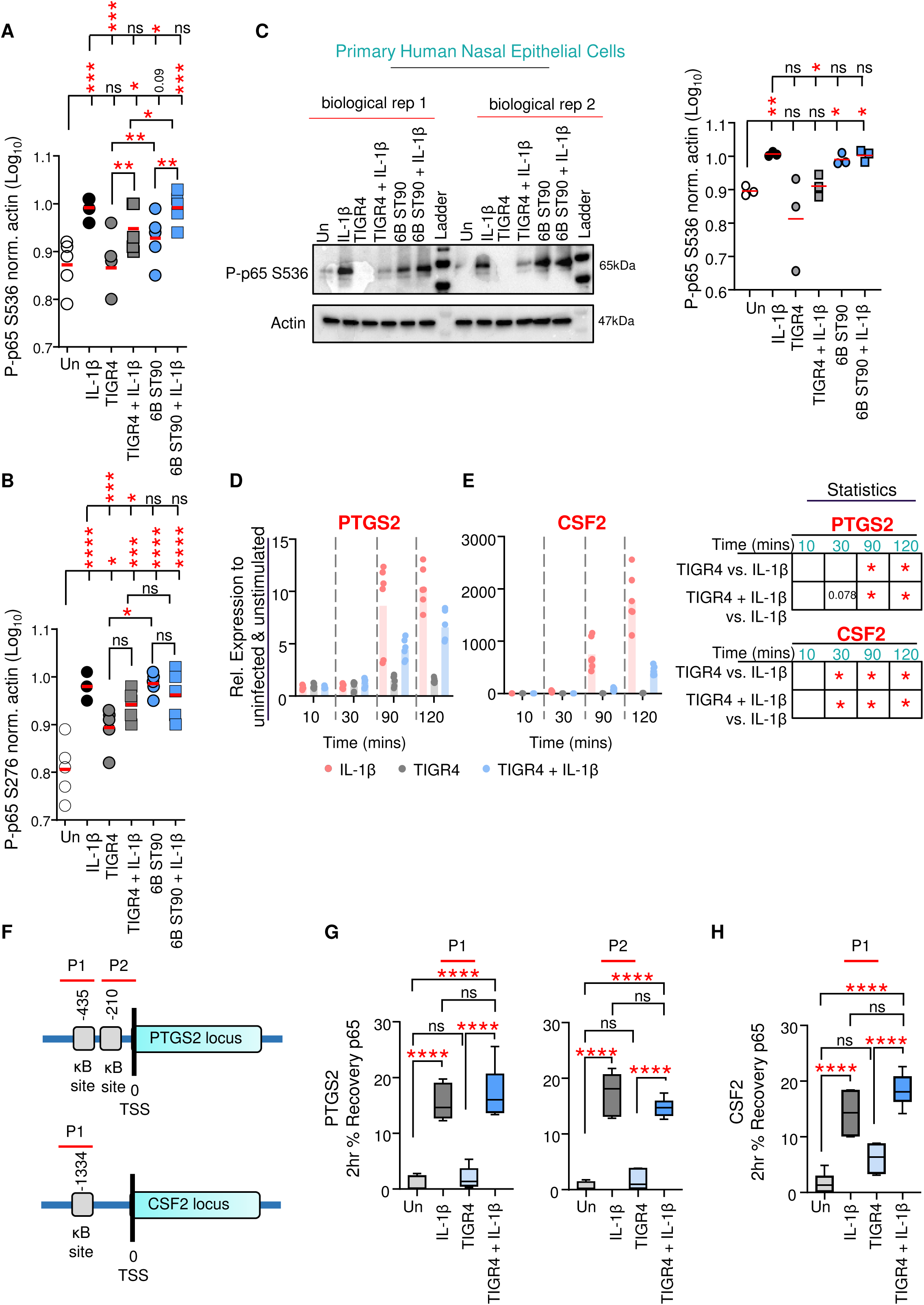
TIGR4 drives disparity in NF-kB gene expression and chromatin binding. A-B) phosphorylated p65 at Serine 536 or Serine 276 to actin. C) Immunoblot of primary nasal whole cell lysates 2 hrs post-challenge with either IL-1β (10 ng/ml), TIGR4 (MOI 20) or 6B ST90 (MOI 20) (± IL-1β; 10 ng/ml) probed for phosphorylated p65 at Serine 536 and actin (n=3 biological). A – C) Dot blot with mean (red line). One-way ANOVA with Dunnett’s multiple comparison post-hoc test. Intergroup comparison paired T-Test. **P ≤ 0.01, ***P ≤ 0.001, ****P ≤ 0.0001. Full blots provided in Supplementary Information 1. D-E) Inflammatory gene transcription profiles by RT-qPCR for PTGS2 & CSF2 of A549 cells over a 2 hr time course with either IL-1β or TIGR4 (± IL-1β 10 ng/ml; MOI 20). Graphed as the relative expression of each indicated transcript to matched uninfected/unstimulated control per time point (n = 3 biological replicates; 2 technicals per replicate). Displayed as a dot plot with each data point and a bar representing the mean. Statistics table reporting significance calculated by unpaired T.Test at each time point indicated. * = significant; full details of analysis in Sup. Table 4. Schematic representation of PTGS2 or CSF2 promoter with ChIP-qPCR primer locations and NF-κB sites denoted as predicted by Alibaba2-curated eukaryotic transcription factor DNA-binding motifs from the TRANSFAC database ^68^. G-H) 10µg chromatin input from A549 cells either untreated (light gray), IL-1β treated (10 ng/ml; dark gray) or 2 hrs post-challenge with TIGR4 (± IL-1β; 10 ng/ml; MOI 20; light blue or dark blue respectively). ChIP-qPCR represented as % recovery against input of p65 at indicated NF-κB sites. Tukey box and whisker plot with defined box boundaries being the upper and lower interquartile range (IQR), ‘whiskers’ (fences) being ± 1.5 times IQR and the median depicted by the middle solid line (n=3 biological replicates with 2 or 3 technicals per replicate). One-way ANOVA comparing all means with Tukey’s multiple comparison post-hoc test. ns=not significant, ****P ≤ 0.0001.

**Sup. Figure 3:**
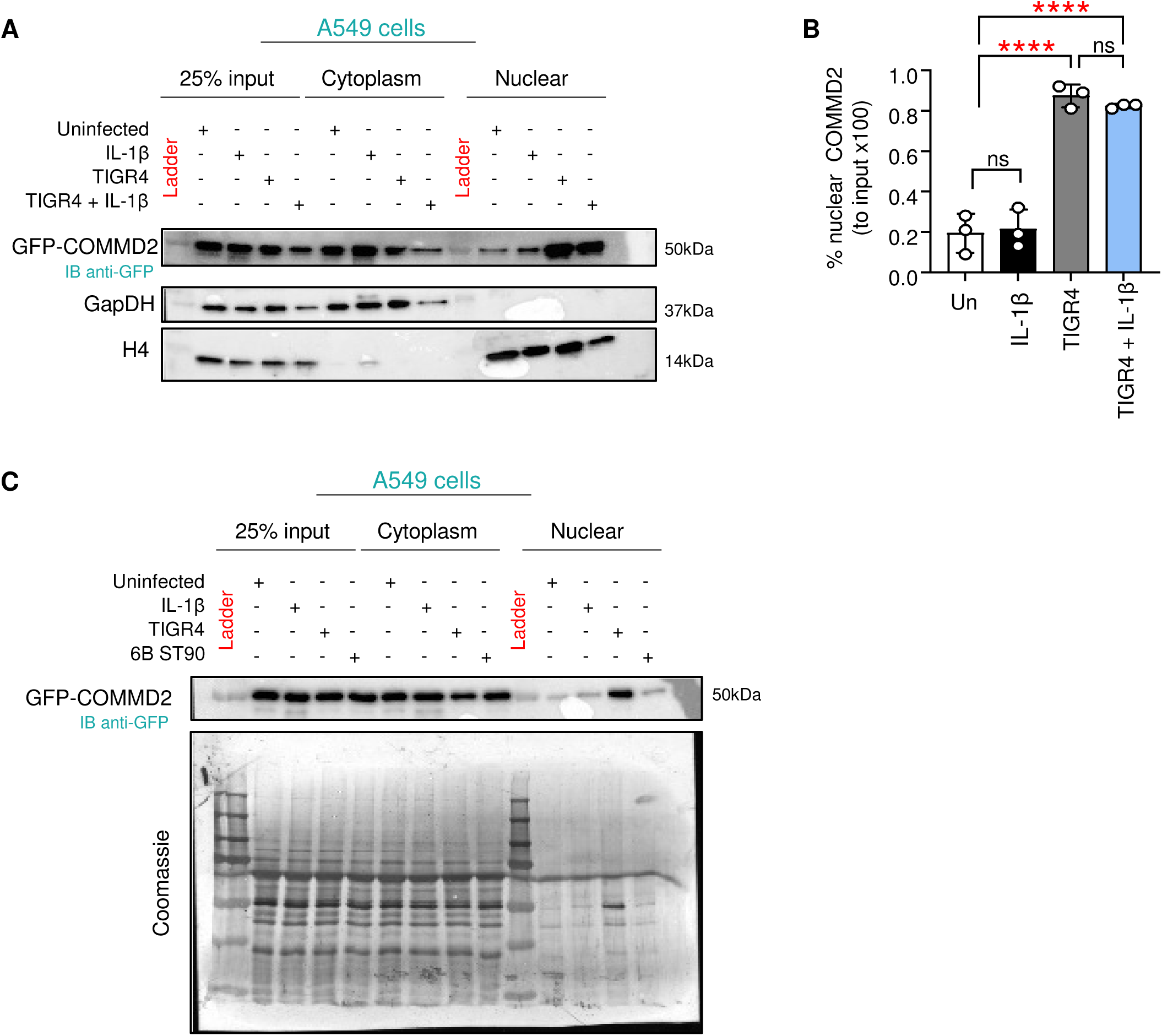
Proteasomal degradation is not involved in TIGR4 mediated p65 turnover. Cell fractions from a stable A549 GFP-COMMD2 cell line 2 hrs post-challenge with either IL-1β (10 ng/ml), TIGR4 (MOI 20) or 6B ST90 (MOI 20). A) Representative immunoblot of cell fractions probed for GFP (COMMD2) enrichment across cellular compartments with GapDH (cytoplasm control) or H4 (nuclear control). B) Quantified nuclear COMMD2 (n = 3 biological). Bar graph with dots representing experiments. One-way ANOVA with Tukey’s multiple comparison post-hoc test. ns = not significant, ****P ≤ 0.0001. C) Representative immunoblot of cell fractions and coomassie stained PVDF membrane.

**Sup. Figure 4:**
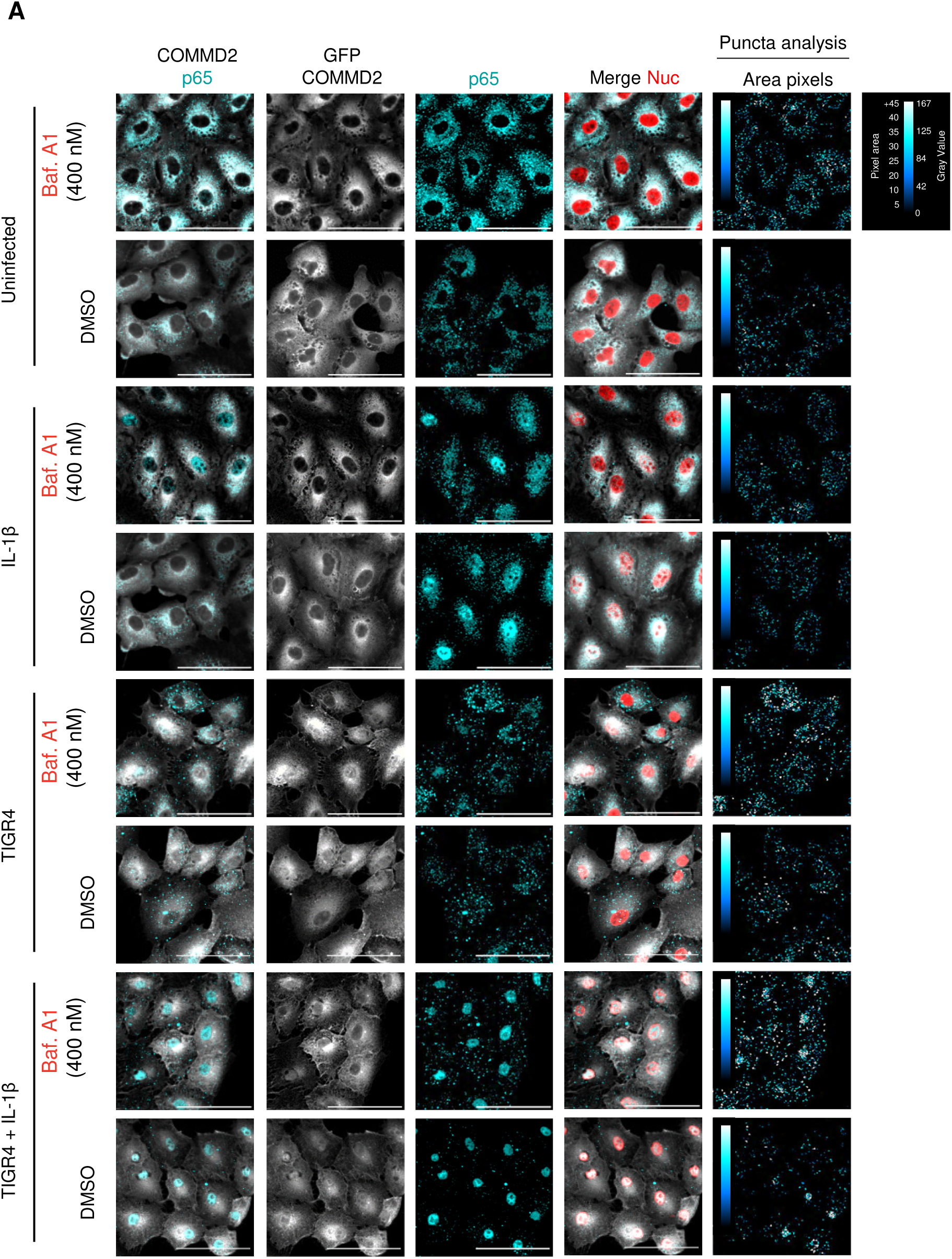
Representative images of COMMD2 & p65 with Bafilomycin A1. Representative confocal microscopy images. Puncta analysis of area in pixels highlighted with color scale.

**Sup. Figure 5:**
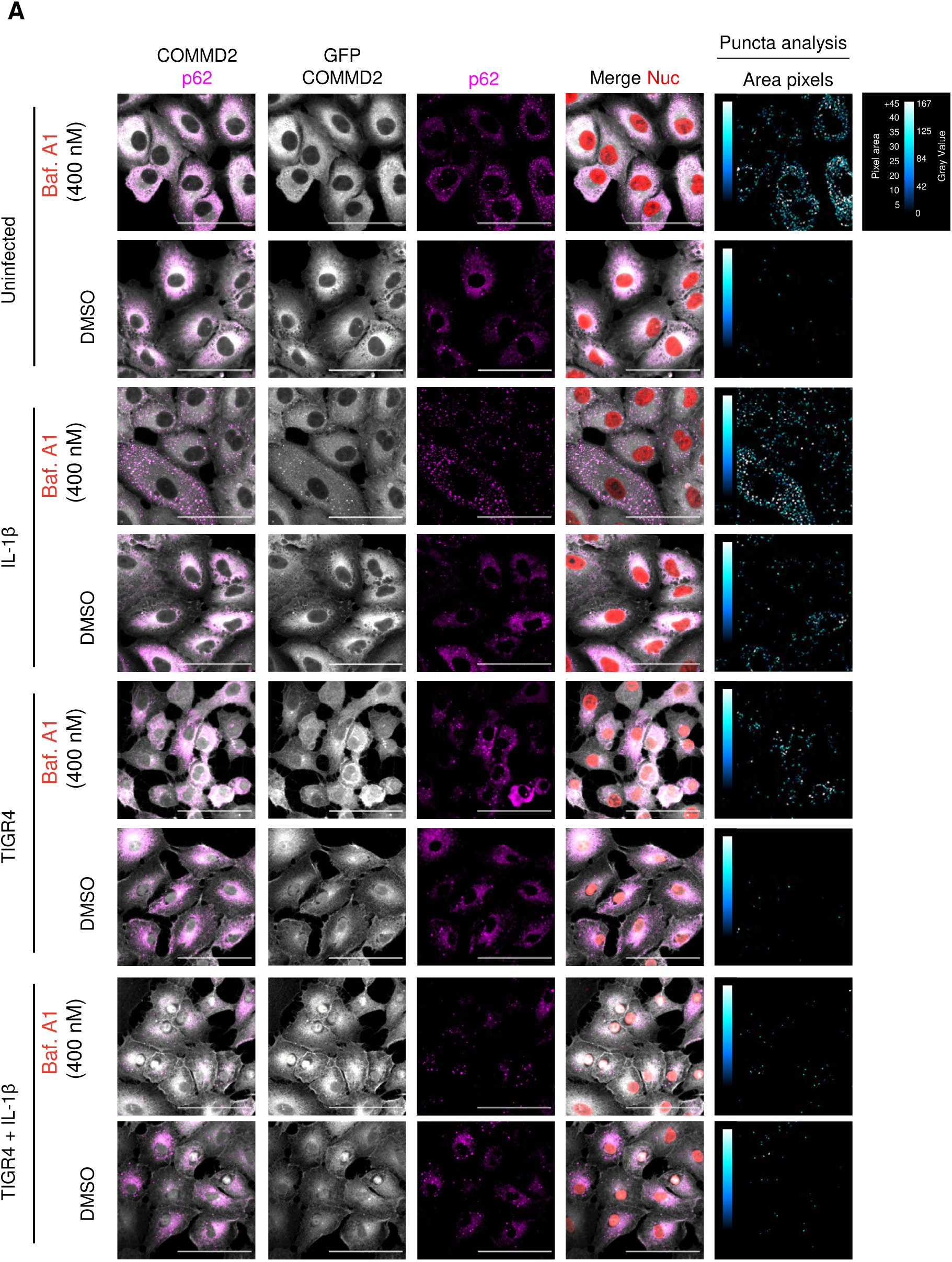
Representative images of COMMD2 & p62 with Bafilomycin A1. Representative confocal microscopy images. Puncta analysis of area in pixels highlighted with color scale.

**Sup. Figure 6:**
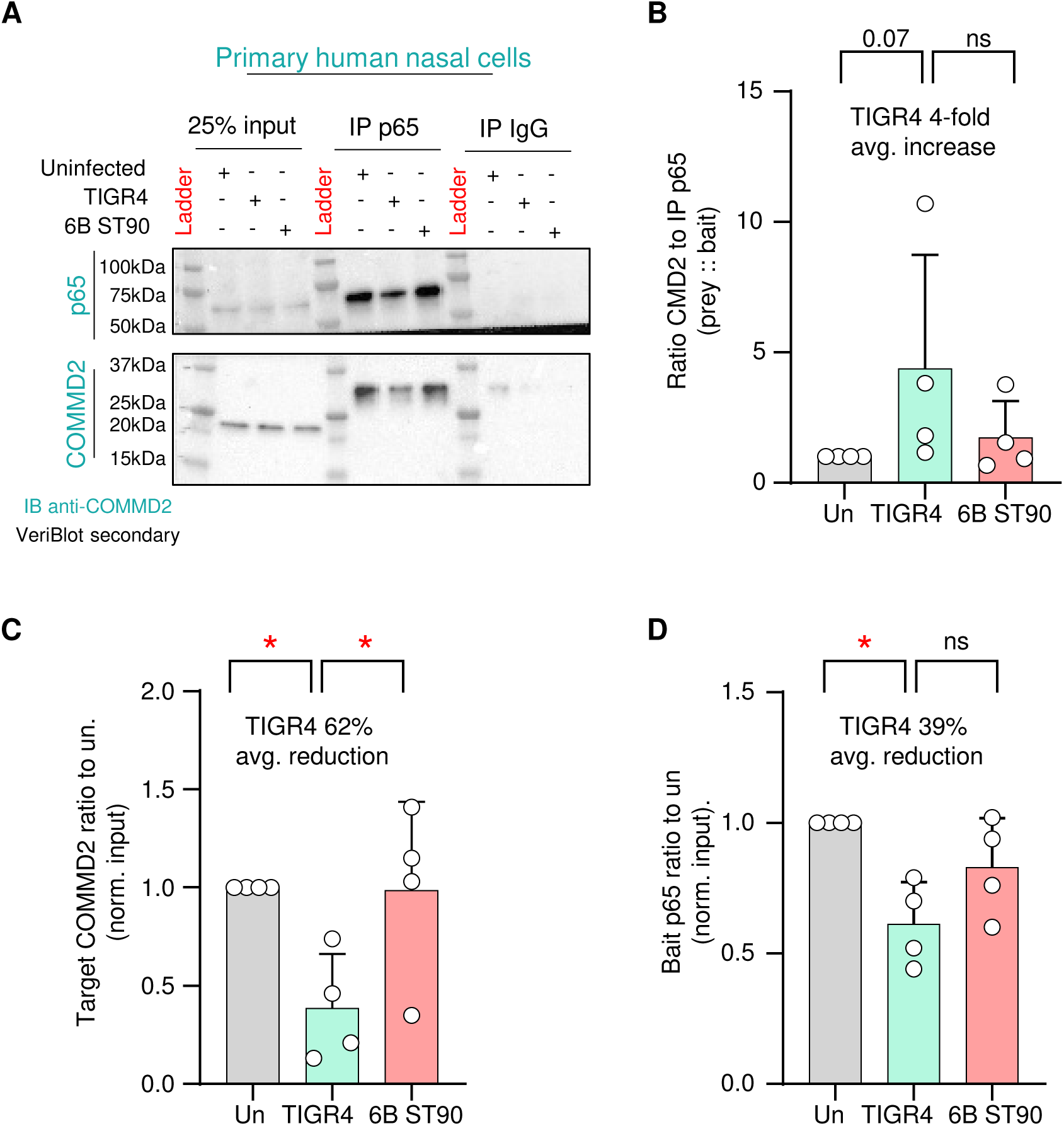
TIGR4 induces endogenous COMMD2-p65 complex. A) Representative immunoblot of p65 immunoprecipitation from primary nasal cells 2 hr post-challenge with either TIGR4 or 6B ST90 (MOI 20; n= 4 biological). B-D) Quantified ratio of p65 to COMMD2 (prey :: bait) normalized to uninfected (B), target COMMD2 ratio to uninfected normalized to input (C) and bait p65 ratio to uninfected normalized to input (D). Dot and bar graph. (B) Kruskal-Wallis test with Dunn’s multiple comparison test. C-D) One-way ANOVA with Bonferroni’s multiple comparison post-hoc test. *P ≤ 0.05.

**Sup. Figure 7:**
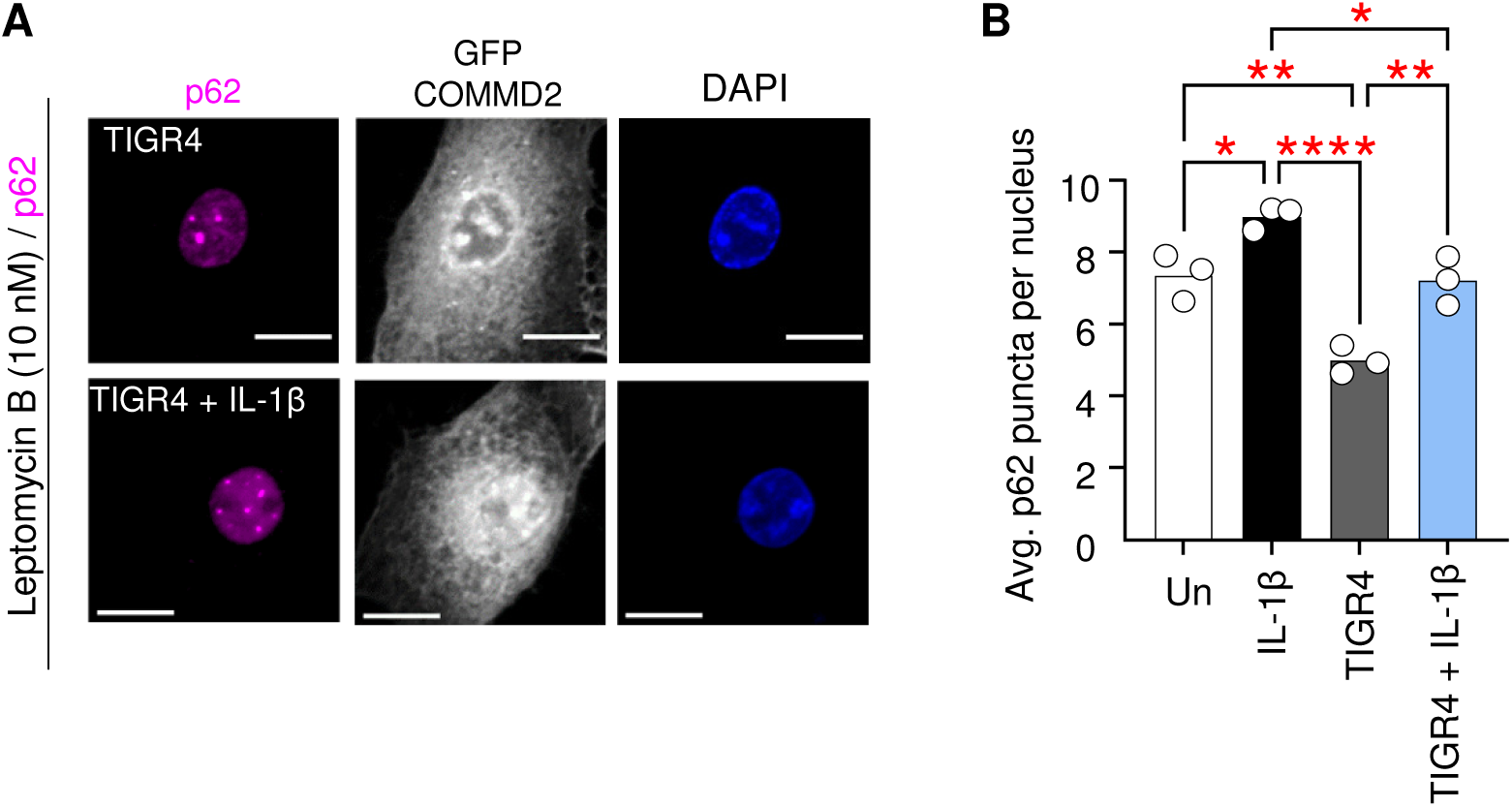
CRM1 inhibition increases p62 puncta. A) Representative immunofluorescence confocal microscopy of stable A549 GFP-COMMD2 pretreated for 3 hrs with Leptomycin B (10 nM) prior to 2 hr challenge with either IL-1β (10 ng/ml) or TIGR4 ± IL-1β (MOI 20). Paraformaldehyde fixed cells stained for p62 (magenta) against GFP-COMMD2 (gray) and nuclei (DAPI; n=3 biological replicates; nuclei counts for Uninfected n=1041, IL-1β n=831, TIGR4 n=1164, TIGR4 + IL-1β n=1269). Scale bar = 10 µm. B) Nuclear p62 puncta average counts per cell. Graphed as mean with dots representing individual experiments. One-way ANOVA comparing all means with Tukey’s multiple comparison post-hoc test. ns=not significant, *P ≤ 0.05, **P ≤ 0.01, ****P ≤ 0.0001.

**Sup. Figure 8:**
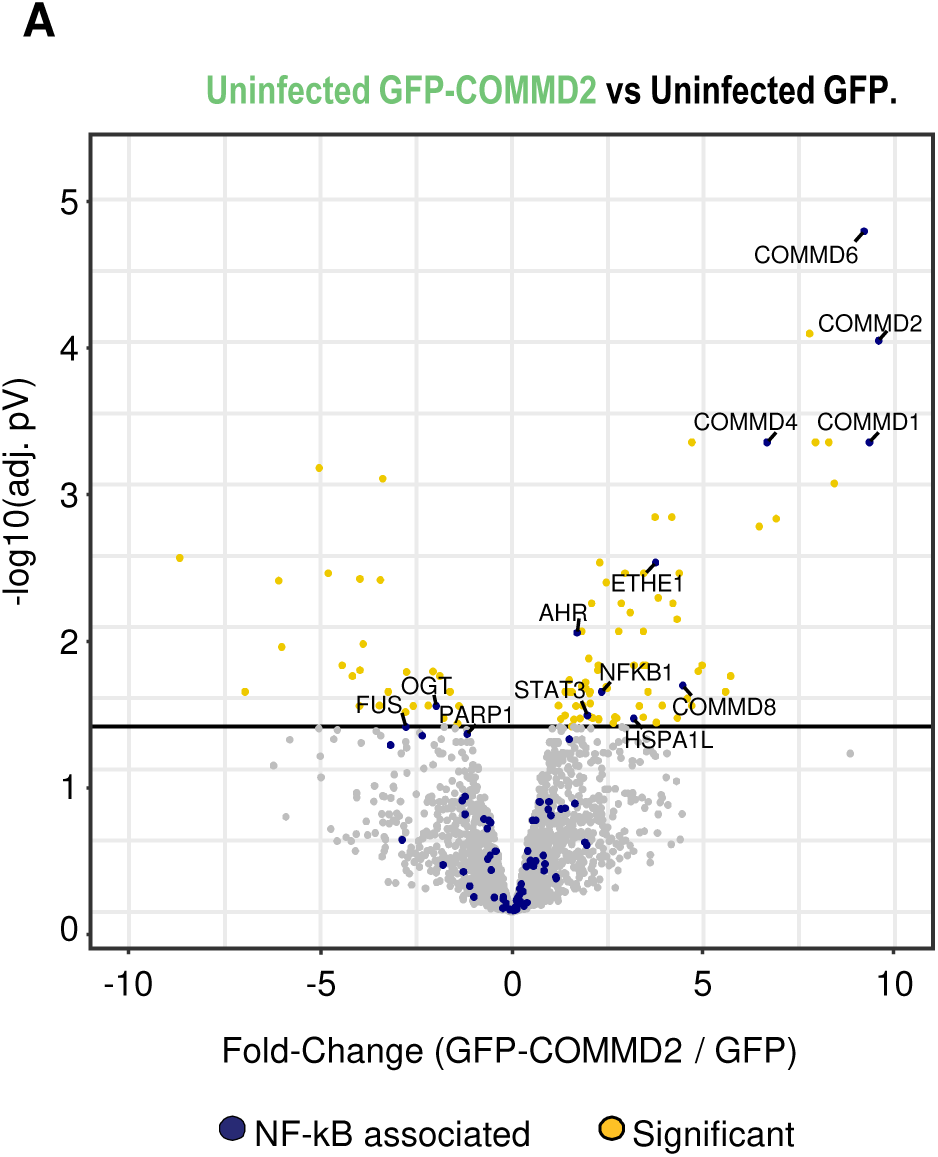
GFP-COMMD2 vs. GFP volcano plot. Mass-spectrometry interactome (n=2 biological replicates per condition) of immunoprecipitated GFP-COMMD2 from a stable A549 GFP-COMMD2 cell line (1×10^7^ cells total) or GFP A549 cell line control (1×10^7^ cells total). Volcano plot of GFP-COMMD2 vs. GFP alone control. Identified interacting partners with known NF-kB p65 interaction in blue and significant targets in yellow. Lines represent −log10(pV) and fold-change cutoffs with targets of interested denoted.

**Sup. Figure 9:**
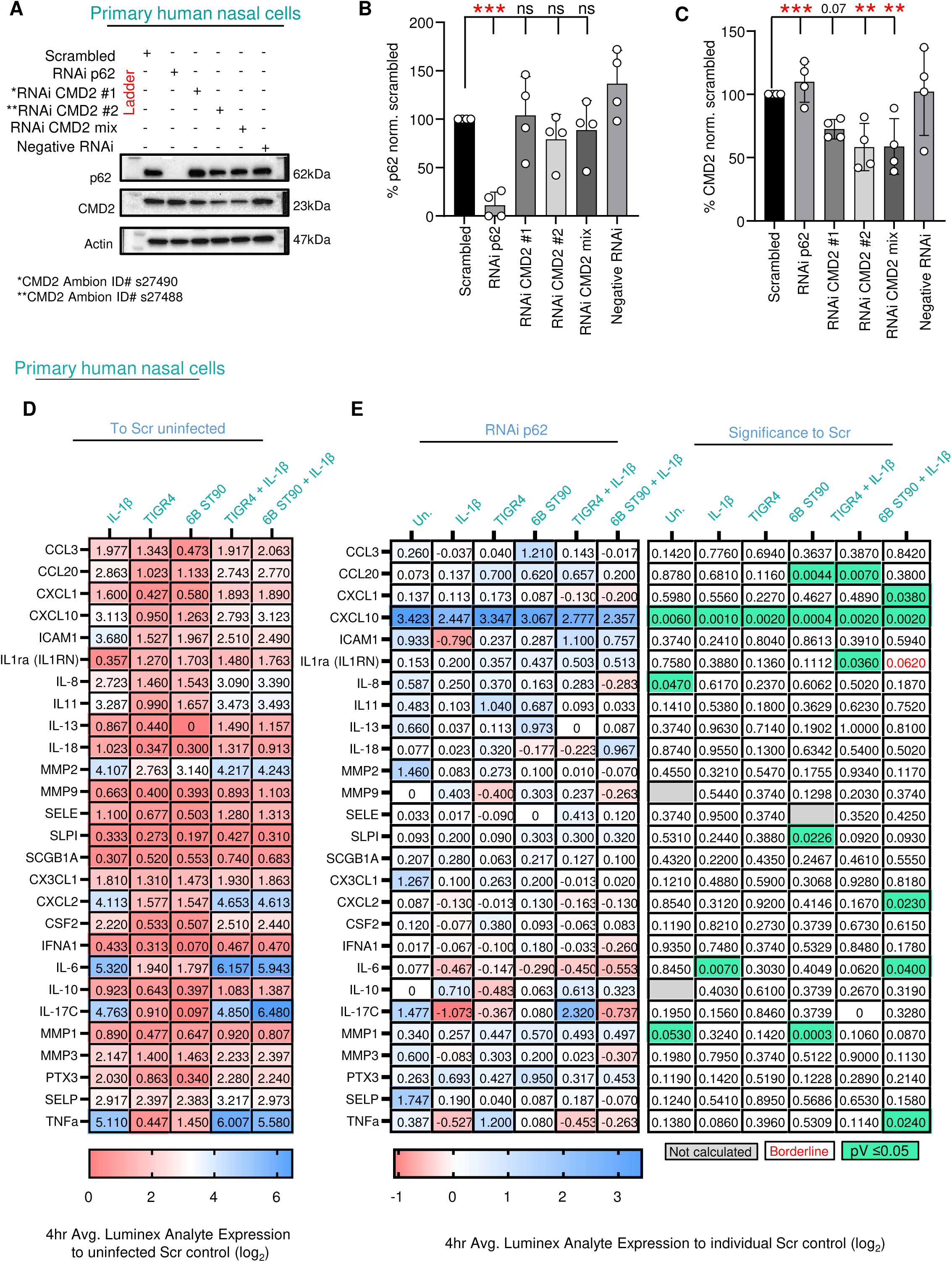
RNAi efficiency in primary nasal cells and Luminex analyte analysis of p62 RNAi. A) Representative immunoblot of RNAi conditions targeting COMMD2 or p62 in primary human nasal epithelial cells from whole cell lysates at 48 hrs of RNAi. PVDF membrane probed for either COMMD2 or p62 protein levels. B & C) Quantification of COMMD2 and p62 protein levels post RNAi. Dot blot with mean (bars) ± STD. One-way matched ANOVA with Dunnett’s multiple comparison post-hoc test comparing RNAi conditions to scrambled control. ns = not significant, **P ≤ 0.01, ***P ≤ 0.001. D) Secreted analytes (Luminex) at 4 hrs from Scr control in primary nasal cells (n=3 biological replicates) challenged with either IL-1β (10 ng/ml), TIGR4 (MOI 20) ± IL-1β or 6B ST90 (MOI 20) ± IL-1β. E) Luminex of secreted analytes at 4 hrs from RNAi p62 in primary nasal cells normalized to scrambled controls (n=3 biological replicates). Cells were challenged with either IL-1β (10 ng/ml), TIGR4 (MOI 20) ± IL-1β or 6B ST90 (MOI 20) ± IL-1β. Individual p values shown in comparison to respected Scr control. Calculated by unpaired T. Test. Full details of analysis in Sup. Table 4.

**Sup. Figure 10:**
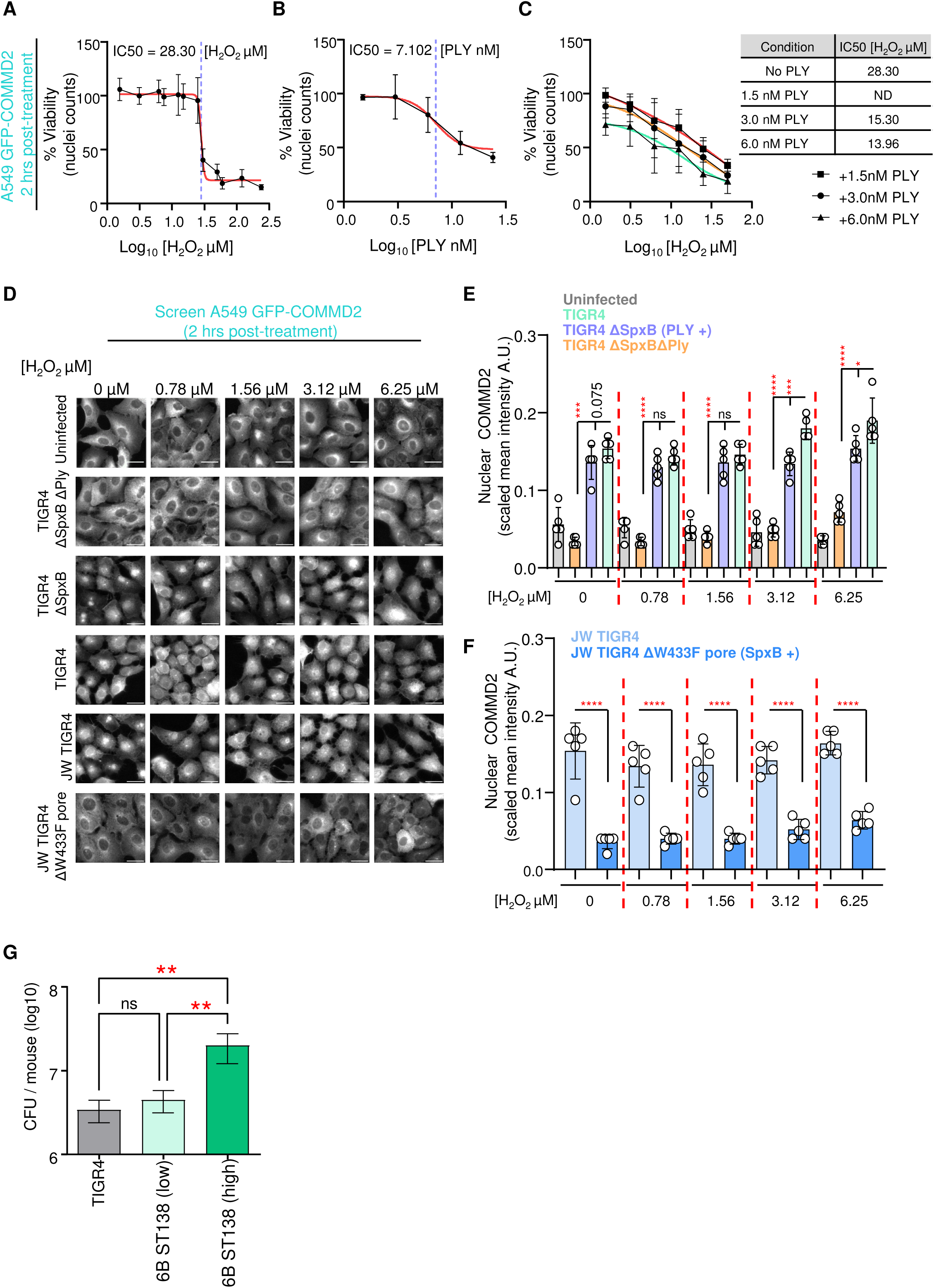
CMD2 nuclear translocation screen with pneumococcal mutants and hydrogen peroxide. A-C) Titration of hydrogen peroxide (A), purified Pneumolysin toxin (B), or their combination (C) upon viability of the GFP-COMMD2 A549 stable cell line (n=4 biological replicates). Graphed as a dot ± STD of biological replicates for IC_50_ calculations by 4 parameter variable slope curve (red) with IC_50_ denoted in purple dashed line. Tabular comparison of IC_50_. D) Representative deconvolution epifluorescence microscopy of A549 GFP-COMMD2 (gray) cells 2 hr post-challenge with either MOI 20 for each bacterial strain JW TIGR4 (wildtype), JW TIGR4 Δ*ply* W433F, TIGR4 (wildtype), TIGR4 *ΔspxB* or TIGR4 *ΔspxBΔply* under titrated hydrogen peroxide treatment (0 – 6.25µM). E-F) Graphed as Dot blot with mean (bars) ± STD. One-way matched ANOVA comparing selected columns with Bonferroni’s multiple comparison post-hoc test. ns = not significant, *P ≤ 0.05, **P ≤ 0.01, ***P ≤ 0.001, ****P ≤ 0.0001. G) Inocula of intranasal mouse challenge shown as bar graph of mean ± STD. One-way ANOVA comparing all means with Tukey’s multiple comparison post-hoc test. ns=not significant, **P ≤ 0.01.

